# A comparison of feature selection methodologies and learning algorithms in the development of a DNA methylation-based telomere length estimator

**DOI:** 10.1101/2022.04.02.486242

**Authors:** Trevor Doherty, Emma Dempster, Eilis Hannon, Jonathan Mill, Richie Poulton, David Corcoran, Karen Sugden, Ben Williams, Avshalom Caspi, Terrie E Moffitt, Sarah Jane Delany, Therese M. Murphy

**Author notes:** These authors contributed equally (Sarah Jane Delany, Therese M. Murphy).

## Abstract

The field of epigenomics holds great promise in understanding and treating disease with advances in machine learning (ML) and artificial intelligence being vitally important in this pursuit. Increasingly, research now utilises DNA methylation measures at cytosine-guanine dinucleotides (CpG) to detect disease and estimate biological traits such as aging. Given the high dimensionality of DNA methylation data, feature-selection techniques are commonly employed to reduce dimensionality and identify the most important subset of features. In this study, we test and compare a range of feature-selection methods and ML algorithms in the development of a novel DNA methylation-based telomere length (TL) estimator. We found that principal component analysis in advance of elastic net regression led to the overall best performing estimator when evaluated using a nested cross-validation analysis and two independent test cohorts. In contrast, the baseline model of elastic net regression with no prior feature reduction stage performed worst - suggesting a prior feature-selection stage may have important utility. The variance in performance across tested approaches shows that estimators are sensitive to data set heterogeneity and the development of an optimal DNA methylation-based estimator should benefit from the robust methodological approach used in this study. Additionally, we observed that different DNA methylation-based TL estimators, which have few common CpGs, are associated with many of the same biological entities. Moreover, our methodology which utilises a range of feature-selection approaches and ML algorithms could be applied to other biological markers and disease phenotypes, to examine their relationship with DNA methylation and predictive value.

## 1. Background

Epigenetic biomarkers such as those derived from 5-methyl cytosine (DNA methylation) can help to address important questions across a myriad of biological fields. DNA methylation-based estimators and classifiers are models developed using statistical and machine learning (ML) methods that utilise DNA methylation data for the estimation of a range of variables including biological age, telomere length (TL), disease diagnosis, smoking status and body mass index (BMI). As illustration of their utility - instead of using chronological age, an imperfect surrogate measure of the ageing process, DNA methylation-based age estimates that indicate biological aging of a person can be used to investigate the impact of stress factors on individuals of the same chronological age [1]. Similarly, many studies utilise self-reported smoking information whose inaccuracies can propagate through medical research studies [2, 3]. An accurate epigenetic estimator of smoking history can serve to ameliorate this issue. Furthermore, the importance of epigenetic mechanisms such as DNA methylation has become evident in the pathogenesis of various diseases, with DNA methylation markers emerging as potential clinical biomarkers [4]. Recently, an expanding body of work involving DNA methylation-based ML approaches and their utility to estimate and predict a range of quantitative traits including chronological age [5-14], epigenetic smoking scores [15-17] and body mass index (BMI) [18] have increasingly become apparent.

Current limitations in the accuracy and efficacy of developed estimators include the fact that each estimator’s performance is data set dependent, leading to variability in the markers selected across different estimators of the same traits. Furthermore, studies that utilise DNA methylation-based data for purposes such as disease classification or estimation of a trait sometimes only explore a single feature-selection method or, at best, a relatively limited range. Data sets generated from high-throughput DNA methylation arrays measure methylation levels at CpG sites along the DNA sequence, thus providing features for statistical and ML models. They typically contain extremely high numbers of features in combination with small sample sizes, and can suffer from the curse of dimensionality [19]. Feature-selection methods can return a subset of variables that may reduce the effects of noise or irrelevant features while still providing useful prediction results [20]. Additionally, they can reduce computation time and potentially improve predictive performance [21, 22]. They are often employed to reduce the high dimensionality of input datasets, mitigate collinearity, and, in conjunction with a learning algorithm, estimate the quantitative trait of interest.

There are different approaches to feature-selection. Filter methods, a commonly used approach for high dimensional data, usually perform feature ranking based on statistical or information theoretic measures which generate a score that captures the amount of information each independent variable has about the dependent variable [22]. These fast methods are independent of the predictive model but can suffer from selecting redundant features. Choosing the best filter approach depends very much on the level of computational resources available to the researcher [22]. Wrapper methods in contrast utilize performance metrics of the predictive model to select the best feature subsets while embedded methods include feature-selection in the process of the modelling algorithm’s execution [23, 24].

These feature-selection methods naturally lead to dimension reduction but there are other methods which can also achieve this. Principal component analysis (PCA) is the dominant feature transformation technique, capturing most of the variance in the data by projecting the data into a reduced feature space. This transformative method, which is commonly used in ML, yields a set of orthogonal variables (principal components) that are linear combinations of the original variables [25]. In this sense, it has less utility for identifying specific features e.g., for acquisition of an explicit biological signature. However, its ability to tackle multicollinearity (the presence of strong relationships between variables in a data set), which can impact the performance of statistical and ML-based models, makes it a potentially powerful technique for development of an estimator primarily sought for its predictive capability [26].

Using telomere length as a vehicle to explore a robust methodology, our study wishes to build on previous ML studies in the epigenomics field by evaluating a range of feature-selection methods and ML algorithms to identify a novel DNA methylation-based estimator of TL and investigate its association with other health-related demographics. Telomere length - DNA repeat structures which are located at the ends of each chromosome and have a crucial role in maintaining genomic stability - has emerged as a promising biomarker for biological age [27]. In recent years an association between TL and epigenetic processes has been hypothesised. Recently, Lu et al. [28] developed a DNA methylation-based estimator of TL (DNAmTL) which utilised 140 cytosine-phosphate-guanine dinucleotides (CpGs) using a regression-based ML approach (i.e. elastic net [29]), highlighting the power of ML methods to develop robust DNA methylation-based estimators.

In this study, we first reviewed the literature to ascertain feature-selection and learning algorithms commonly used in the epigenomics field for the estimation of quantitative traits. Due to the popularity in the published literature of elastic net penalised regression for epigenetic aging signatures and its use in the previously reported DNA methylation-based TL estimator [28], this approach forms the baseline algorithm in our study. Many studies also utilise some form of initial feature-selection in advance of applying elastic net, therefore we also investigated applying a range of feature-selection methods as an initial step. Previous studies have utilised association tests corrected for multiple testing, using false discovery rate (FDR) thresholds [6-8]. The FDR has demonstrated ability to detect true positives while controlling Type I errors at a designated level with methods such as the Benjamini and Hochberg step-up procedure (BH) [30] and q-value [31] arguably the most widely used and cited approaches for FDR control in practice [32]. In addition to association tests, studies have employed thresholding of correlation between CpGs and the quantitative target feature using Pearson’s correlation coefficient [6, 13, 33-35], and mutual information [36, 37].

Additionally, ensemble ML algorithms such as random forest can be used to obtain a set of ranked features and is among the methods investigated in this study. Several comparative studies demonstrated that Support Vector Regression (SVR) performed well as an alternative to elastic net and other regression-based approaches [8, 35, 38], and we also explore this method along with several other regression algorithms in conjunction with feature-selection. In total, we develop and evaluate nine TL estimators, each adopting a different feature-selection methodology. In addition to identifying a novel DNA methylation-based estimator of TL, we aim to develop a robust methodology utilising ML algorithms which could be applied to other biological markers and disease phenotypes, to examine their relationship with DNA methylation.

## 2. Methods

### 2.1 Data description

The data is comprised of 3 cohorts (Dunedin, EXTEND and TWIN), which have measures of both TL and Illumina DNA methylation array data (summarised in Table 1). The Dunedin data set pertains to a cohort (n=1037) born in Dunedin, New Zealand between April 1972 and March 1973 as described elsewhere [39]. Assessments occurred at a range of ages, most recently at age 45 when 938 of the living 999 study members took part. We have used data from two sweeps of the study i.e., ages 26 and 38. With two time points, most participants contributed two samples and, after preprocessing, 1631 samples were utilised. The DNA methylation data was derived from whole blood and measured using the Illumina Infinium HumanMethylation450 BeadChip [40] (Illumina, CA, USA). TL was measured using a validated quantitative PCR (qPCR) method [41], as previously described [17].

**Table 1:** Summary Details - Data Sets

| Name | Samples | Features | Target Type |
| --- | --- | --- | --- |
| Dunedin | 1631 | 431553 | Continuous |
| EXTEND | 192 | 430574 | Continuous |
| TWIN | 178 | 482369 | Continuous |

The validation EXTEND data set (n=192) is a subset of the Exeter 10,000 epidemiological cohort as described in detail elsewhere [42]. The second validation TWIN data set (n=178) involves a cohort previously recruited from a multi-centre collaborative project aimed at identifying DNA methylation differences in MZ twin pairs discordant for schizophrenia as described elsewhere [43]. The same laboratory measured TL by a validated qPCR method [44] in both the EXTEND and TWIN validation cohorts. DNA methylation levels were measured in the same labs using the Illumina Infinium methylation array platform for both the EXTEND and TWIN validation cohorts. For all 3 data sets the methylumi package [45] was used to extract signal intensities for each CpG probe and perform initial quality control, with data normalization and pre-processing using the wateRmelon package as described previously [46]. All experiments were performed in accordance with national guidelines and regulations as well as the Declaration of Helsinki. The Dunedin study participants gave written informed consent, and study protocols were approved by the NZ-HDEC (Health and Disability Ethics Committee). For both the EXTEND and TWIN studies, informed consent was obtained from participants and ethical approval was obtained from University of Exeter Medical School Research Ethics Board.

DNA methylation levels at CpG sites formed the input features for models while the target variable was TL. Relative telomere lengths in the Dunedin, EXTEND and TWIN datasets were adjusted based on plate ID via linear regression to control for plate-to-plate variation. Any CpG columns containing missing values were not utilised in the development of the final estimators, with the initial set of CpG sites restricted to 417,690 common to all three data sets.

### 2.2 Modelling overview

For the application of the regression algorithms and the feature-selection methods used in this paper, the *Python* software package *scikit-learn* (version 0.23.2) was utilised [47]. Stage 1 of our analysis involved the comparison of a range of models that utilised different feature-selection methods in conjunction with elastic net regression i.e., DNA methylation-based TL estimators. A nested cross-validation process, which includes hyperparameter tuning, was utilised for model comparison which has been suggested as a more appropriate approach when assessing a model, providing a reliable estimate of the true error [48, 49], and avoiding information leakage from test data into the training process. A range of performance metrics are reported in this study. These include the Mean Absolute Error (MAE), Mean Absolute Percentage Error (MAPE), and Pearson correlation coefficient between predicted and actual TL (described in Section 1 of Supplementary Information (S.I.)).

The nested cross-validation (CV) process to assess model performance was employed on the Dunedin data set. The data was first split into train and test data sets in 80%:20% proportions respectively i.e., DS_train_ and DS_test_. Each feature-selection method (or feature transformation method in the case of PCA) was applied wholly to DS_train_, which yielded a subset of features. Using only these features, 3-fold cross-validation was then applied to DS_train_ using elastic net in order to choose the best model hyperparameters (via a random search) based on the mean absolute error (MAE) metric. The next step involved constructing an elastic net model on DS_train_ (using the discovered feature subset and best parameters). This model (i.e., the estimator) was then used to predict the instances contained in DS_test_, yielding a range of pertinent performance scores (MAE, MAPE and correlation coefficient between predicted and actual TL) to indicate estimator efficacy. This constitutes one iteration of the 5-fold outer cross-validation that is part of the 5 × 3 nested CV process, and it is therefore repeated for each of the other outer cross-validation iterations. The nested CV process was conducted for each of the 9 investigated estimators (the baseline using elastic net and 8 feature selection/transformation methods followed by elastic net). The estimators were then compared by considering the combination of the aforementioned performance scores. The feature-selection methods and their application in this first stage of the analysis are outlined in Section 2.3.2.

The next stage of the analysis (Stage 2) involved using the same 9 approaches from Stage 1 to construct DNA methylation-based estimators from the full Dunedin data set. These estimators were then tested on the two sets of independent data (EXTEND and TWIN). Unlike the performance results from the first stage which were generated from nested CV analysis on a single data set, the results of this second stage of analysis represent estimator performance on data originating from different laboratories and populations from which the estimators were trained. As such, this extended our model comparison process from nested CV on a single data set to evaluation on 2 independent data sets.

To obtain feature subsets from the Dunedin data, the 9 approaches were applied to the full Dunedin data set, in each case returning an *N* feature subset. With these *N* features, 10-fold cross-validation, with a grid search, was conducted on the Dunedin data to find a good set of model hyperparameters. Using the *N* features and best identified parameters, estimators were then constructed from the Dunedin data and used to make predictions on the 2 independent validation sets. The majority of subjects in the Dunedin data set contributed 2 samples for analysis. Therefore, all data partitioning (such as within the cross-validation process) was implemented such that, where 2 samples pertain to a subject, both samples were contained within the same partition. This prevents information leakage, ensuring subject independence across data partitions.

### 2.3 Elastic net regression and feature-selection methods

#### 2.3.1 Elastic net regression

Given its common usage in DNA methylation-based age prediction studies [5, 7, 8, 10] and its application in a recent TL estimator [28], elastic net regression was used as a baseline model for TL estimation from DNA methylation data in this study. Regularized models, like elastic net regression, facilitate the selection of predictive CpGs among correlated markers when the ratio of features to samples is very large [50]. With too much freedom, a model can be prone to over-fitting due to too many available features. One solution to this is regularisation where extra terms are introduced into the objective function that penalise extreme values of regression coefficients and, thus, encourages them to take small values unless absolutely necessary. Elastic net regression is an embedded feature-selection approach, and its algorithm includes dimensionality reduction or intrinsic feature-selection. This may be a reason for its popularity in this domain.

In elastic net regression, the input variables, *X*, and output variable, *y*, are represented by a least squares relationship, given as: y = β_0_ + **βX** + **e**, where β_0_ is the intercept, and **β** and **e** are the vectors of regression coefficients and residuals respectively. Elastic net employs a mixture of both the l_1_ and l_2_ penalties and can be represented as:

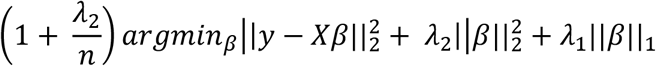

where 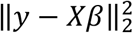 is the l_2_-norm loss function (i.e., the residual sum of squares), 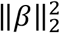 is the l_2_-norm penalty on β, λ_2_ is a regularisation (complexity) parameter, ‖β‖_1_ is the l_1_-norm penalty on β and λ_1_ is a regularisation parameter. Setting 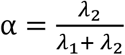, the elastic net estimator is shown to be equivalent to the minimisation of: 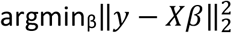, subject to the penalty 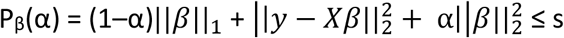, for some s [51]. The parameter α is known as the mixing parameter as it defines how the l_1_ and l_2_ regularisation is mixed. The elastic net model was tuned using the regularisation parameters, λ, and the mixing parameter α. Over a grid search of parameter values, the parameter pairing which returned the minimum MAE corresponded to the optimal model from the cross-validation parameter tuning process. The parameter ranges used in the grid search process were [1 × 10^−5^, 1 × 10^−4^, 1 × 10^−3^, 1 × 10-^2^, 1 × 10^−1^, 0.25, 0.5, 1, 10, 100] and [0.01, 0.1, 0.3, 0.5, 0.7, 0.9, 0.95, 0.99, 1] for the λ and α parameters respectively.

#### 2.3.2 Feature-selection methods

The feature-selection methods used in this work include a number of filter feature-selection approaches which typically require some user-specified threshold that determines the set of features used. For the F-test approach, the false discovery rate is set at a threshold value - we use both 0.05 and 0.01 as these are typically used in this domain. Utilising these thresholds will return a set of features.

The other filter feature-selection approaches used provide a ranking of the features. Using Pearson and mutual information, the correlation strengths and actual information gain between each independent variable and the dependent variable respectively can be ranked. With support vector regression, the regression coefficients can be ranked, while the decision tree ensemble approach (random forest) returns a feature importance score that allows feature ranking.

The reduced feature set can be selected using a threshold of some sort on this ranking, e.g., by choosing those with scores higher than a threshold score or by specifying a number of features to use. We select the number of features to use for each method by determining the set that performs best on the training data. We chose to investigate models with successively larger feature subset sizes, in an effort to find the model that corresponded to the error minima/correlation maxima. In Stage 1 of the analysis (nested CV), elastic net regression models were created after initial feature selection using each of the four ranking feature-selection methods, i.e., for ranked feature subsets of 50, 100, 150, 200, 250, and in steps of 250 thereafter up to 20000 features. Nested CV performance scores were then plotted against feature subset size to yield curves for examination of the effect of increasing feature subset size, and from which an optimal point (feature subset size) could be selected for the stage 2 and stage 3 analyses.

The other feature-selection approaches used in this study include the embedded feature-selection techniques of elastic net, which is our baseline approach, and gradient boosting which is a decision tree ensemble technique which includes regularisation. Both these techniques yield a unique feature subset from the application of the algorithm. We also used PCA which is a feature transformation approach. The principal components were derived from training data and used to project both training and test data into the training PCA subspace. More detail is given on the individual feature selection/transformation methods in Table 2, while Table 3 outlines commonly used feature-selection approaches from a range of studies that utilise DNA methylation data.

**Table 2:** Summary of feature-selection techniques used in this study.

| Type | Feature-Selection Method | Description |
| --- | --- | --- |
| Filter - Univariate | Correlation-based (Pearson's $r$) | Pearson's correlation for two random variables $a$ and $b$ is given as: $\rho(a,b)=\text{cov}(a,b)/(\sigma_a\sigma_b)$ where $\text{cov}(a,b)$ is the covariance or cross-correlation of $a$ and $b$ and $\sigma$ denotes the standard deviation [52]. Pearson's correlation coefficient ($r$) is a measure of linear association, and it is used in epigenetic studies to measure the strength of association between each feature and the response variable. A threshold value is usually applied, excluding all features below that value. |
|  | Multiple hypothesis testing (F-test with FDR) | F-test statistic or associated p-value can be used as a threshold score for association between a feature and response. The false discovery rate (FDR) can be used to detect true positives while controlling Type I errors at a designated level. F-tests between the methylation value of each CpG and TL are conducted, with those CpG sites with FDR (Benjamini-Hochberg [30]) less than a specified value being selected. |
| | Mutual Information | Mutual information can be formulated as: $MI = H(x) + H(x\vert y)$, where $H(x)$ is the entropy of feature $x$ and $H(x\vert y)$ denotes the entropy of feature $x$ after observing feature $y$. The values of mutual information per feature are typically ranked with a threshold utilised to remove the most redundant features (CpGs) [37]. |
| Filter - Regression | Support Vector Regression | The absolute value of the weights (coefficients) yielded by the support vector regression (SVR) algorithm can be utilised to create a set of ranked features. In the case of a linear kernel, the SVR model can take the form: $\text{prediction}(x) = b + \mathbf{w}^T \mathbf{x}$ where $\mathbf{w} = \sum_i \alpha_i x_i$, with the vector of weights $\mathbf{w}$ directly accessible. The features with higher absolute weights are considered to be more likely to be useful for model training and prediction, conversely smaller weights are thought to not have a large influence on predictions [53]. |
| Filter - Ensemble | Random Forest Regression | Random forests [54] can handle correlated data and high dimensionality [55]. This ensemble method for classification and regression utilises bagging (subsets of samples) and boosting (subsets of features) to ensure diversity across constituent tree models. As trees only use a portion of samples in their construction, the remaining samples can be used to generate feature importance scores via feature value shuffling, with the impact of this shuffling assessed over the whole ensemble [56]. |
| Embedded | Elastic net | This is a regularised regression method and embedded feature-selection approach. It includes both the $l_1$ and $l_2$ norms in the |
|  |  | objective function and tunes the bias towards one of the norms using a hyperparameter [57]. |
|  | XGBoost | XGBoost [58] utilises gradient boosted decision trees and can generate feature importance scores through the degree that each feature split point enhances performance, weighted by the number of observations relating to a node [59]. BoostARoota [60] is an embedded method which uses XGBoost as its base learner and returns a reduced feature set through regularisation. |
| Transformative | Principal Component Analysis (PCA) | PCA is applied to a data set containing variables, which are, in general, inter-correlated. It finds new variables which are linear combinations of the original variables that maximise variance but are uncorrelated with each other. [61]. |

**Table 3:**
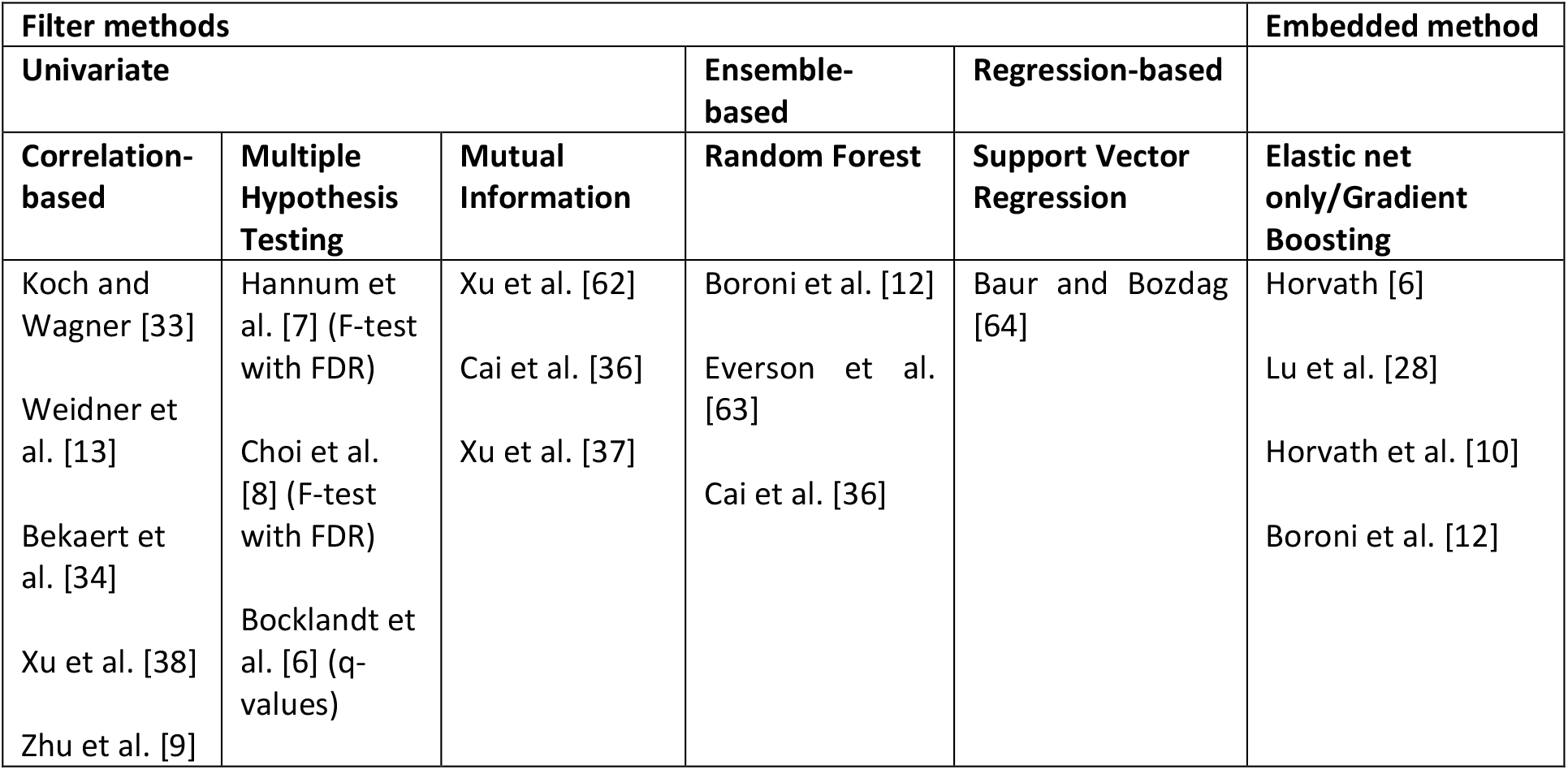
Commonly used feature-selection approaches in DNA methylation-based studies.

### 2.4 Other regression-based learning algorithms

In the third stage of analysis (Stage 3), estimators were constructed by combining a range of different regression algorithms with the feature subset utilised in the estimator judged to be the most promising from the Stage 2 analysis. These included partial least squares regression, multi-layer perceptron, least angle regression and support vector regression – these models had relatively low computational overhead, unlike some other algorithms which would require long run times, given that feature set sizes can still be relatively large after the feature-selection stage. The same training, feature-selection, cross-validation and testing methodology was used as described for the Stage 2 analysis.

### 2.5 Statistical analysis

Pearson’s correlation coefficient was used to assess the strength of linear association between actual and estimated TL in all cohorts. In addition, Pearson’s correlation was calculated between the three TL measures (actual TL, MI-EN TL and PCA-EN TL) and age in the EXTEND cohort. The strength of correlation (Pearson) between age acceleration (obtained by regressing DNAmAge [5] on age) and both actual and predicted TL was also assessed.

To avoid age confounding potential relationships between actual and estimated TL measures and age-related traits, age-adjusted measures were generated for the EXTEND data. This was achieved by regressing actual TL, MI-EN TL and PCA-EN TL on age, with the raw residuals of this process being defined as TLadjAge, MI-ENadjAge and PCA-ENadjAge respectively. Accordingly, Pearson’s correlation was assessed between these age-adjusted measures and estimated blood cell counts for the EXTEND data set. Furthermore, repeated measures correlation was used to examine the correlation between actual TL/MI-EN TL/PCA-EN TL and actual blood cell counts available for the Dunedin cohort. This method accounts for repeated measures for the same individuals at ages 26 and 38 – addressing non-independence among observations by statistically adjusting for inter-individual variability using analysis of covariance [65].

Multiple linear regression models were used to explore a range of biological correlates i.e., the association between variables such as TL (both actual and estimated) and participant traits (e.g., age and sex), while controlling for confounders. Tests were two-sided and the statistical significance level was defined as p < 0.05.

The estimator previously developed by Lu et al. (DNAmTL) [28] was compared with the estimators constructed in our study. The ability of the compared estimators was assessed via MAE, MAPE, and the Pearson correlation coefficient between predicted and actual TL, on both our independent data sets (EXTEND and TWIN).

## 3. Results and discussion

### 3.1 Nested cross-validation analysis on Dunedin data set

A comparison of nested CV MAE on the Dunedin data set for the 9 feature-selection models investigated in this study is shown in Figure 1. Models utilise elastic net regression following application of each feature-selection/transformation method, with the exception of the baseline model which applies elastic net regression with no prior feature-selection stage.

**Figure 1:**
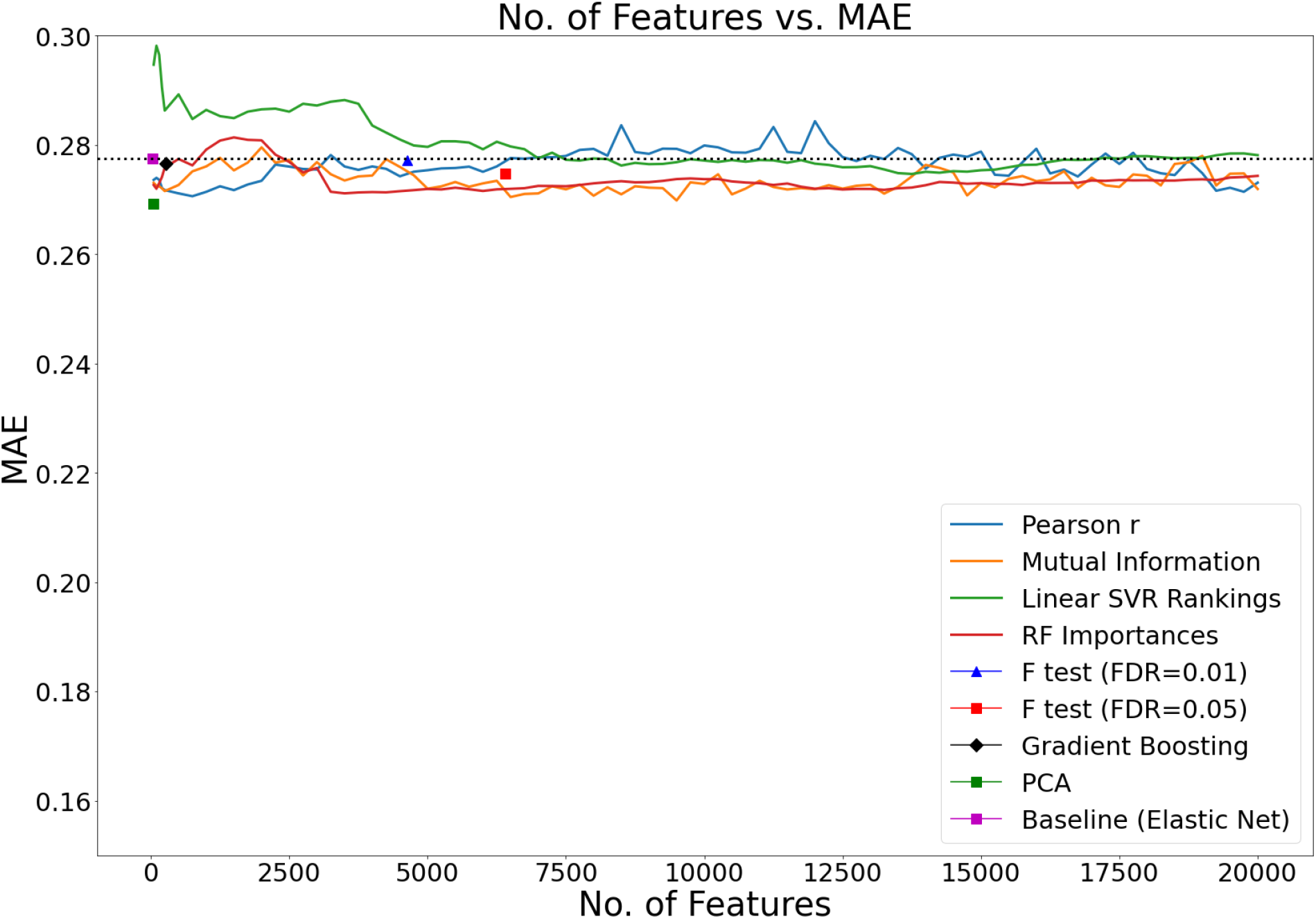
Comparison of 9 feature selection/transformation models using the MAE of the 5 × 3 nested CV analysis. The four line plots relate to those feature-selection methods that yield an explicit feature ranking e.g., mutual information, thus allowing plotting of successively larger feature subsets (models were constructed for feature sets of sizes 50, 100, 150, 250 and every 250 features thereafter up to 20000 – each of these ranked feature sets were later passed to the elastic net regression stage for further feature selection). These line plots can be assessed to identify potentially useful feature subset sizes e.g., at error minima. The five other feature-selection methods (points on the graph) are those which yield a single feature subset, either by being given a specified threshold value (such as for F-tests with FDR) or being the result of an embedded method (gradient boosting and the baseline model (elastic net)). The number of features shown for these five methods represents the final feature set size after application of elastic net regression. The dotted black line represents the baseline performance.

Figure 1 shows that the model that utilises the Pearson correlation-based feature rankings performed relatively well at a lower number of features. On closer inspection of the graph, a feature set size of approximately 750 is close to the minimum error observed for that model. We will utilise this feature set size in stage 2 of our analysis where models are constructed based on the full Dunedin data set and tested on the 2 independent data cohorts. The model that utilises linear SVR feature rankings can be seen to progressively improve up to approximately 8500 features and arguably plateaus thereafter, while the model that implements PCA in advance of elastic net regression, yields the lowest error across all models. Although PCA is not strictly a feature-selection technique but rather a feature transformation method, we were interested to assess its performance, given that multicollinearity is a known challenge in high-dimensional data when applying statistical methods [26, 66]. The results suggest that the orthogonality of the transformed variables (principal components) mitigated, to some extent, the issue of correlated predictors.

In addition to assessing the MAE, it is useful to also consider the Pearson correlation coefficient between predicted and actual TL and previous studies report this when considering the efficacy of TL estimation [28, 67]. For the same models shown in Figure 1, Figure 2 shows the Pearson correlation coefficient between the predicted and actual TL. In Figure 2, the Pearson correlation-based feature ranking method with elastic net again indicates relatively good performance at a lower feature set size, with a peak observed at approximately 750 features. It is notable that performance is seen to increase for this model as the number of features approaches 20,000. However, it is important to consider that, in the field of epigenetics, there can be an advantage in discovering relatively smaller feature subsets (biological signatures) given that this facilitates both model interpretation and downstream biological interpretation of DNA methylation estimators of biological traits. However, larger feature numbers may have more utility to estimate telomere length in population-based studies.

**Figure 2:**
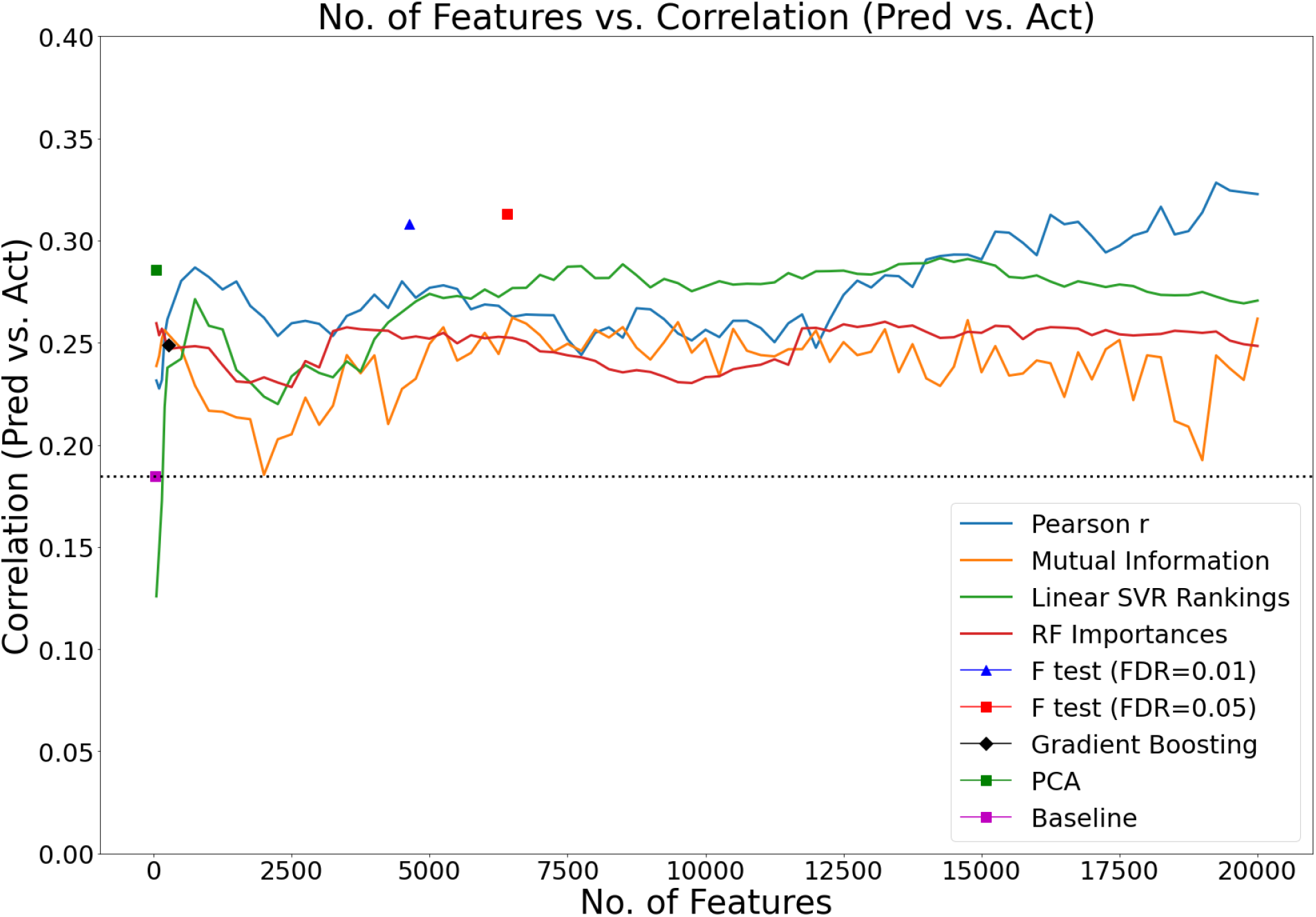
Comparison of the 9 feature-selection models using the Pearson correlation between predicted and actual TL for the 5 × 3 nested CV analysis. These line plots can be assessed to identify potentially useful feature subset sizes e.g., at correlation maxima. The dotted black line represents the baseline model (elastic net with no prior feature selection).

Given that Pearson’s correlation indicates how strongly two variables are linearly related, two methods that stand out as performing well relative to the others are the two F-tests with FDR control - exhibiting higher Pearson’s correlation (*r* > 0.3) between actual and predicted TL (Figure 2). Interestingly, hypothesis testing with FDR control is known to be a popular method in DNA methylation-based age estimation studies. Again, the application of principal component analysis before the elastic net regression stage yields a relatively good result (*r* ≈ 0.3). It is notable that the baseline method of elastic net regression without any prior feature-selection stage performs poorly in this analysis (*r* < 0.2), suggesting that a feature-selection stage in advance of elastic net regression is beneficial and should be explored as part of the estimator discovery process. When seeking an optimal DNA methylation-based estimator, the methods and algorithms used to develop this will naturally vary due to the heterogeneity and diversity of datasets – currently most DNA methylation datasets are too small and not sufficiently representative to yield a general-purpose estimator. This highlights the importance of a robust development methodology, as presented here, in the pursuit of a DNA methylation-based estimator of traits.

The average number of features remaining after both the initial feature selection/transformation and elastic net regression stages for the models constructed using nested CV is shown in Table 4. Additionally, the bar charts in Figure 3 and Figure 4 show the extent of the feature reduction process for each of the 9 tested models. The left y-axes scales are logarithmic due to the large differences in features shown. Consider an example, the Pearson correlation-based feature-selection model was trained with the top ranked 750 features. The decision to use 750 features was based on observation of minima and peaks in Figure 1 and Figure 2 respectively from the nested CV analysis stage i.e., we effectively chose the best model from the models tested over the range 50 to 20000 Pearson correlation ranked features as described in Section 2.3.2. In Figure 3, for the Pearson correlation model, the blue bar indicates the 750 features retained after the initial feature-selection stage, with the adjacent orange bar indicating that approximately 70 features remain after application of the elastic net regression algorithm to the 750-feature data set.

**Table 4:** Features remaining in models after both feature-selection stages for the nested CV analysis (Stage 1). As described in Section 2.3.2, the optimal number of features for the four ranking feature-selection methods (Pearson correlation, mutual information, linear SVR and random forest) were selected from analysis of Figures 1 and 2. For example, the minimum MAE corresponded to passing 3250 features from the random forest feature ranking to the elastic net regression stage, resulting in an average of 406 features being selected. The figures denote the average (rounded) from the five training sets of the nested CV process described in Section 2.2. Values in parentheses denote the standard error of the mean. * denotes principal components.

| **Models developed using nested CV** | **Average features remaining after initial feature-selection stage** | **Average features remaining after elastic net stage (Std. Error)** |
| --- | --- | --- |
| Baseline (Elastic Net) | - | 30(16.3) |
| F-test (FDR: 0.01)/ Elastic Net | 23685 | 4637 (283.5) |
| F-test (FDR: 0.05)/ Elastic Net | 55750 | 6398 (672.2) |
| Gradient Boosting/ Elastic Net | 584 | 264 (8.7) |
| Pearson correlation/ Elastic Net | 750 | 68 (22.1) |
| Mutual Information / Elastic Net | 6500 | 394 (109.6) |
| Linear SVR/ Elastic Net | 8500 | 4453 (12.6) |
| Random Forest Regression/ Elastic Net | 3250 | 406 (10.6) |
| PCA/Elastic Net | 1072* | 41(9.5)* |

**Figure 3:**
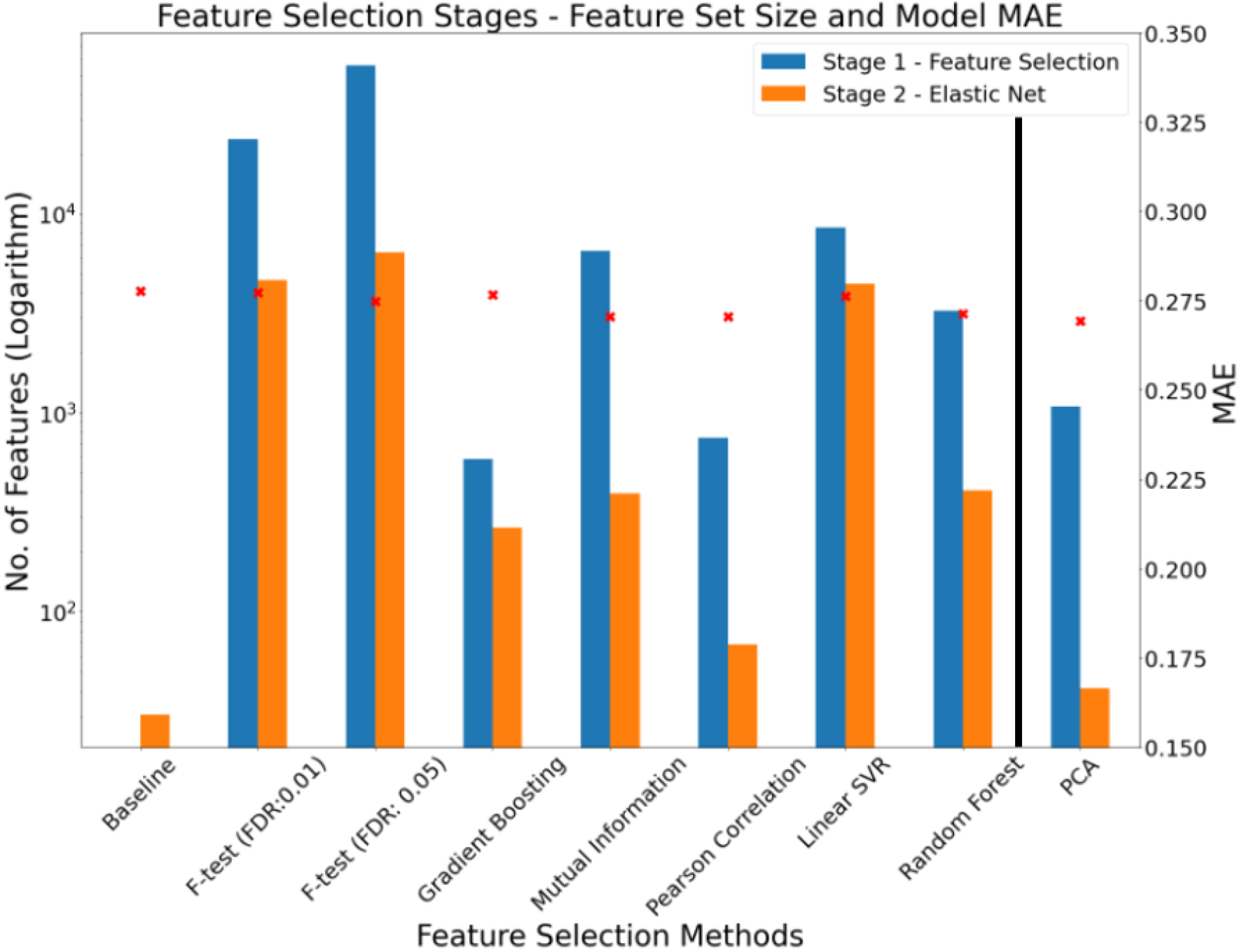
Number of features selected for each model at initial feature-selection stage and elastic net stage. The left axis denotes the number of features (logarithm scaled) with the right axis showing the MAE for each model. The model that utilised PCA in advance of elastic net (to the right of the vertical black line) is shown apart, as PCA is technically a feature transformation method and, as such, the feature count refers to the number of principal components (transformed features). The number of features shown for those models that utilise explicit feature rankings (mutual information, Pearson’s correlation, linear SVR and random forest) pertain, in each case, to the optimal model from all models tested with ranked feature sets in defined steps between 50 and 20000 (as specified in Section 2.3.2).

**Figure 4:**
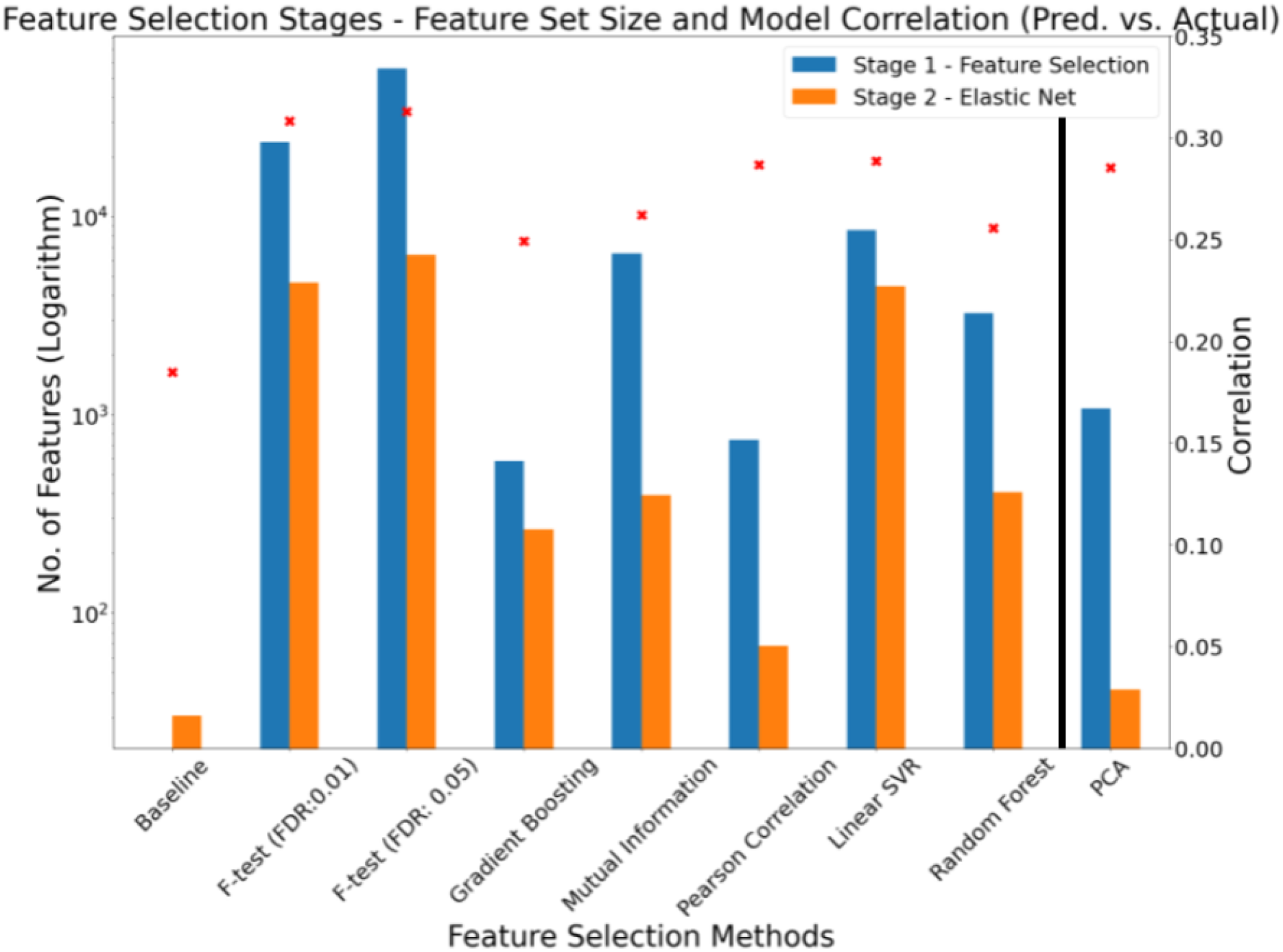
Number of features selected for each model at initial feature-selection stage and elastic net stage. The left axis denotes the number of features (logarithm scaled) with the right axis showing the Pearson correlation between predicted and actual TL for each model.

It is notable that the baseline model drastically reduces the initial 417,690 features down to approximately 30 features (i.e., the average number of features that remained in the models constructed from the 5 training sets of the nested CV process). As with Figure 3, Figure 4 shows the same feature reduction information, however, the right y-axis denotes the Pearson correlation between predicted and actual TL. In general, based on this metric, models that retained relatively lower numbers of features yielded the lowest correlation scores.

To investigate how the number of features presented to the elastic net algorithm relates to the number of features selected by it, we plot this relationship over successively larger feature set sizes for each of the 4 models that utilise explicit feature rankings (Figure 5). Interestingly, models that utilise feature rankings derived from the linear SVR and random forest learning algorithms show an essentially monotonically increasing pattern i.e., in general as the number of features input to the elastic net algorithm increases, so too does the number of features selected by it. In contrast, the Pearson correlation-based and mutual information-based methods display significant fluctuations in the number of features retained by the elastic net algorithm.

**Figure 5:**
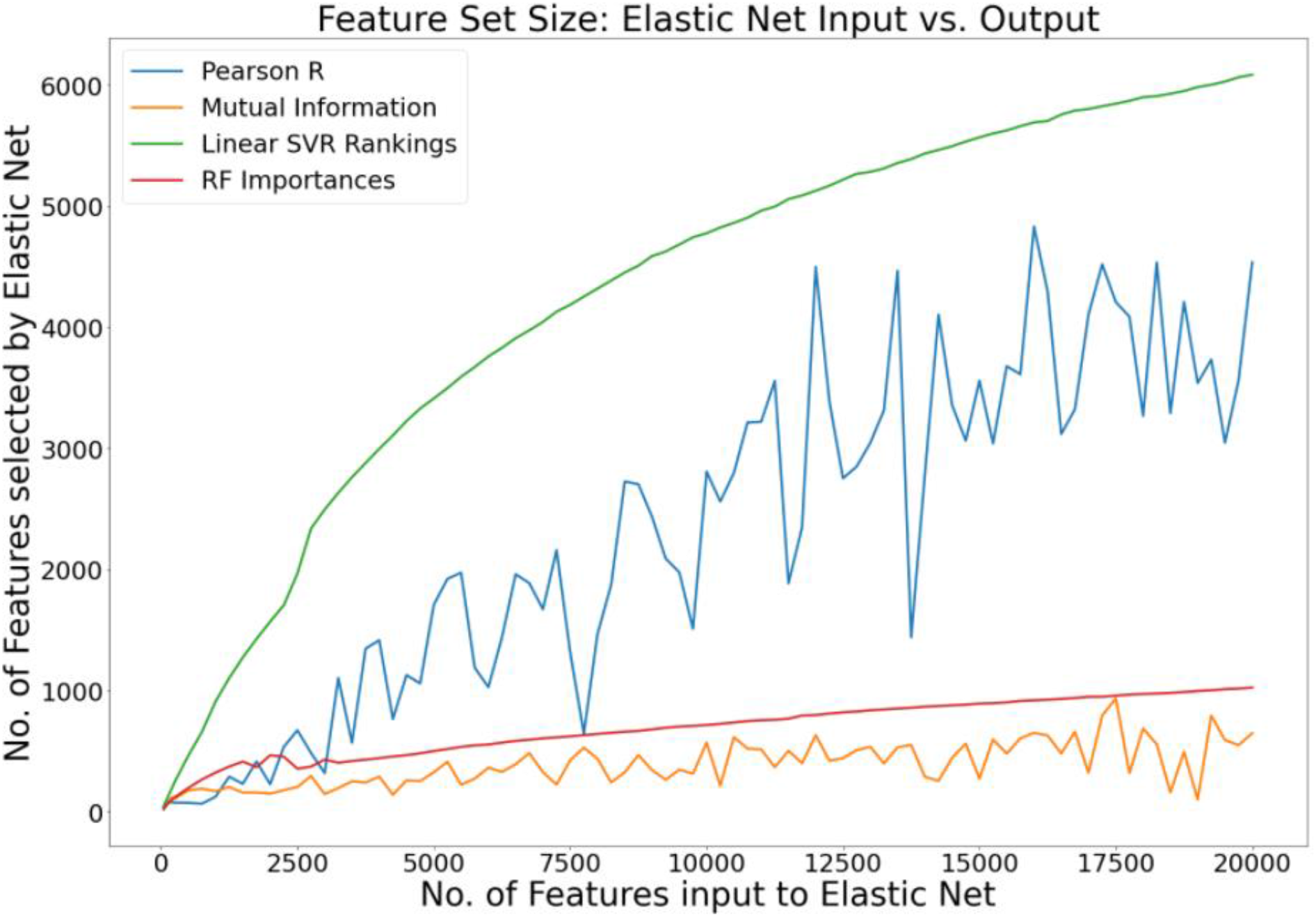
The graph outlines the change in the number of features retained by the elastic net regression algorithm as a function of the input feature set size for the four feature-selection methods that yield explicit feature rankings.

In the case of Pearson correlation, feature subsets of similar size (containing predominantly the same ranked features) passed to the elastic net algorithm can, in some cases, result in substantially different feature set sizes being selected by the elastic net algorithm. The radically different behaviour of the four feature-selection methods and the fluctuating interplay of feature-selection method and regression algorithm further underscores the importance of exploring a range of methods and applying a robust methodology when developing a DNA methylation-based estimator.

### 3.2 Validation on EXTEND and TWIN independent data

Estimators, constructed on the full Dunedin data set, were comprised of specific sets of features after training. These estimators were tested on the 2 independent validation cohorts. Analogous to Table 4, the number of features remaining after both any initial feature-selection stage and the elastic net stage is shown in Table 5.

**Table 5:**
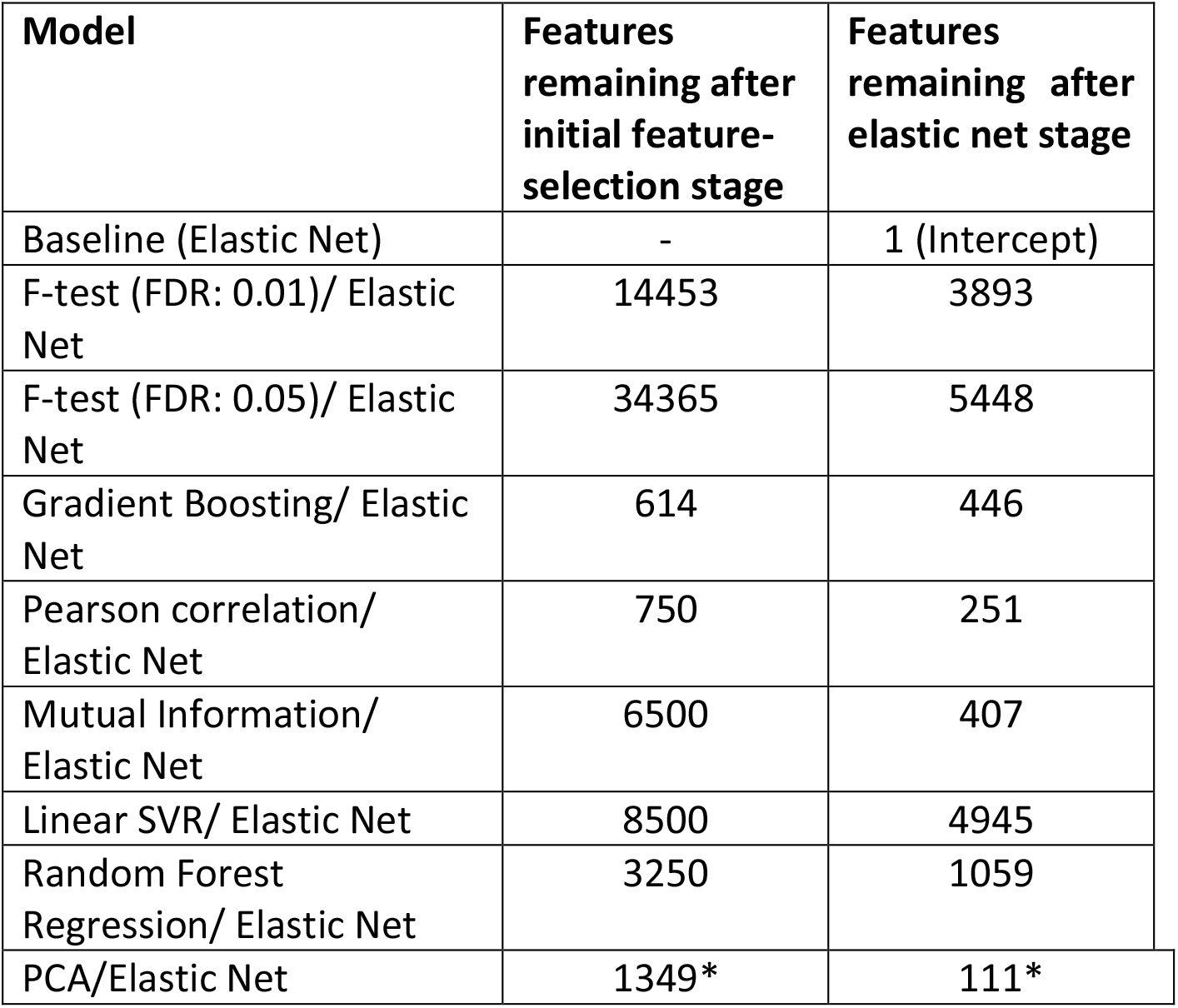
Features remaining after both feature-selection stages for models in the Stage 2 analysis. Note that there is no initial feature-selection stage for the baseline model. * for PCA with elastic net, the number of principal components is shown, as PCA does not return a set of explicit features.

Next, the constructed estimators were tested on both the EXTEND and TWIN data sets. Table 6 denotes the performance measures (MAE, MAPE and Pearson correlation coefficient between predicted and actual TL) of the 9 investigated models, indicating their ability to predict TL in both validation data sets.

**Table 6:** Performance scores for models constructed on the Dunedin data and tested on validation data sets. Metrics include MAE, MAPE and Pearson correlation (predicted and actual TL). The number of features in each estimator is also shown. The estimator *Baseline* refers to the model that utilised elastic net regression with no prior feature-selection stage. Estimator names denote the feature-selection method and the regression algorithm used e.g., F-test-0.01-EN TL refers to the F-test feature-selection stage with FDR of 0.01, followed by application of elastic net. Note that there are no correlation values in the case of the *Baseline* estimator as the model returned the average TL for samples. * denotes principal components.

| Estimator | Feature-Selection Method | Features in estimator (CpGs) | EXTEND |  |  | TWIN |  |  |
| --- | --- | --- | --- | --- | --- | --- | --- | --- |
|  |  |  | MAE | MAPE | Correlation | MAE | MAPE | Correlation |
| **Baseline** | None | 1 | 0.648 | 43.90 | - | 0.756 | 106.99 | - |
| **PCA-EN TL** | PCA | 111* | **0.570** | **39.37** | **0.295** | 0.718 | 112.46 | 0.074 |
| **F-test-0.01-EN TL** | F-test (0.01) | 3893 | 0.614 | 42.93 | -0.003 | **0.728** | **106.52** | **0.119** |
| **F-test-0.05-EN TL** | F-test (0.05) | 5448 | 0.626 | 43.35 | 0.07 | 0.752 | 102.35 | 0.092 |
| **r-EN TL** | Pearson's R | 251 | 0.599 | 41.31 | 0.136 | 0.718 | 116.93 | -0.102 |
| **Boost-EN TL** | Gradient Boosting | 446 | 0.601 | 41.39 | 0.085 | 0.725 | 114.49 | 0.052 |
| **MI-EN TL** | Mutual Information | 407 | 0.640 | 43.51 | **0.203** | 0.753 | 107.83 | -0.067 |
| **LSVR-EN TL** | Linear SVR | 4945 | 0.620 | 43.13 | 0.114 | 0.760 | 108.37 | 0.006 |
| **RF-EN TL** | Random Forest | 1059 | 0.615 | 41.83 | 0.135 | 0.762 | 106.23 | 0.044 |

For the EXTEND data set, the model that applies PCA in advance of elastic net regression (PCA-EN TL) achieves the lowest MAE and MAPE, and the highest correlation between predicted and actual TL. Interestingly, PCA-EN TL was also shown to perform well in the nested CV analysis where models were trained and tested on the Dunedin data set. In this case, the model achieved the lowest MAE (Figure 1) and a relatively high correlation score (Figure 2). Contrastingly, the baseline model (elastic net regression with no prior feature-selection) exhibited poor performance on the data sets utilised in our study. On both independent validation sets, the model returned the average TL when generating predictions for all instances of both data sets. The poor result is somewhat supported by the observed efficacy of the model in the nested CV analysis - Figure 1 shows that the baseline model is among the worst performers in terms of the MAE score, while the model’s estimated TL are the least correlated with actual values (Figure 2). Other notable results include the model that utilised mutual information in advance of elastic net regression (MI-EN TL). This model achieved a correlation of 0.203 between the predicted and actual TL, although the model was seen to perform poorly on the TWIN data set, reporting a small negative correlation (*r* = - 0.067). Of note, no models performed particularly well on the TWIN data.

The use of filter feature-selection techniques in advance of applying elastic net has the disadvantage that user intervention is necessary in most cases. For example, with an F-test, the false discovery rate must be specified, while for the methods that yield explicit feature rankings, the user must explore models with varying feature set sizes to ascertain which model may be optimal. A key aspect of the methodology adopted in our study is that the data used to test performance has not been used for feature-selection – training and testing data must be independent to avoid information leakage from test data into training sets [68].

In contrast, a benefit of utilising embedded methods like elastic net and the gradient boosting implementation used in this study (both without any prior feature-selection step) is that they automatically yield a reduced feature set - which adds to their attractiveness as options. It is important to note that while PCA with elastic net yields the overall best estimator from our tested candidates, the nature of PCA precludes explicit selection of features, yielding instead transformed features that are linear combinations of original features (CpGs). As such, PCA is not commonly used as an initial feature reduction step when developing DNA methylation-based signatures. This limitation, however, is mitigated if the primary motivation for developing an estimator is for predictive purposes where, based on the results of our study, PCA has the potential to yield better estimates of TL than its competitors. A recent study on epigenetic clocks, a widely used aging biomarker derived from DNA methylation data, found that CpG measures can be unreliable due to technical noise. The authors applied PCA in advance of elastic net regression, minimising random noise from CpGs and extracting shared systematic variation in DNA methylation. Technical variance was reduced while preserving relevant biological variance, with the PC-based clocks achieving equivalent or improved prediction of outcomes [69].

Beyond DNA methylation-based estimators of chronological age and TL, studies utilise DNA-methylation data for purposes such as classification of cancer and other diseases [36, 37, 63, 70, 71]. Some studies utilise a single feature-selection technique, while others combine several approaches. Taken together, the robust methodology of comparing and evaluating a range of feature selection and reduction techniques, as demonstrated in our study, could serve to potentially enhance the efficacy and value of DNA methylation-based classifiers and estimators.

### 3.3 Feature-selection with other regression algorithms for estimator development

In addition to elastic net, several other regression algorithms were explored with an initial feature-selection stage. We chose the feature-selection approaches with the best correlation between predicted and actual TL from the validation analysis for both the EXTEND and TWIN data sets. These were mutual information (r=0.203) and F-test (0.01 FDR) (r=0.119) respectively (Table 6). Although the estimator that utilised PCA achieved a better correlation (r=0.295) on the EXTEND data, PCA is not strictly a feature-selection method but rather transforms the original variables into new orthogonal variables. The results of estimators utilising other regression algorithms can be seen in Table 7. Notable results include that the estimator MI-SVR TL obtained a similar correlation between predicted and actual TL (r=0.181) as the MI-EN TL model (Table 6) and that the MI-PLS TL estimator achieves the best MAE and MAPE of all tested estimators on the EXTEND data but a relatively low correlation. However, none of the models that used alternative regression approaches achieved a higher correlation than MI-EN TL for the EXTEND data.

**Table 7:**
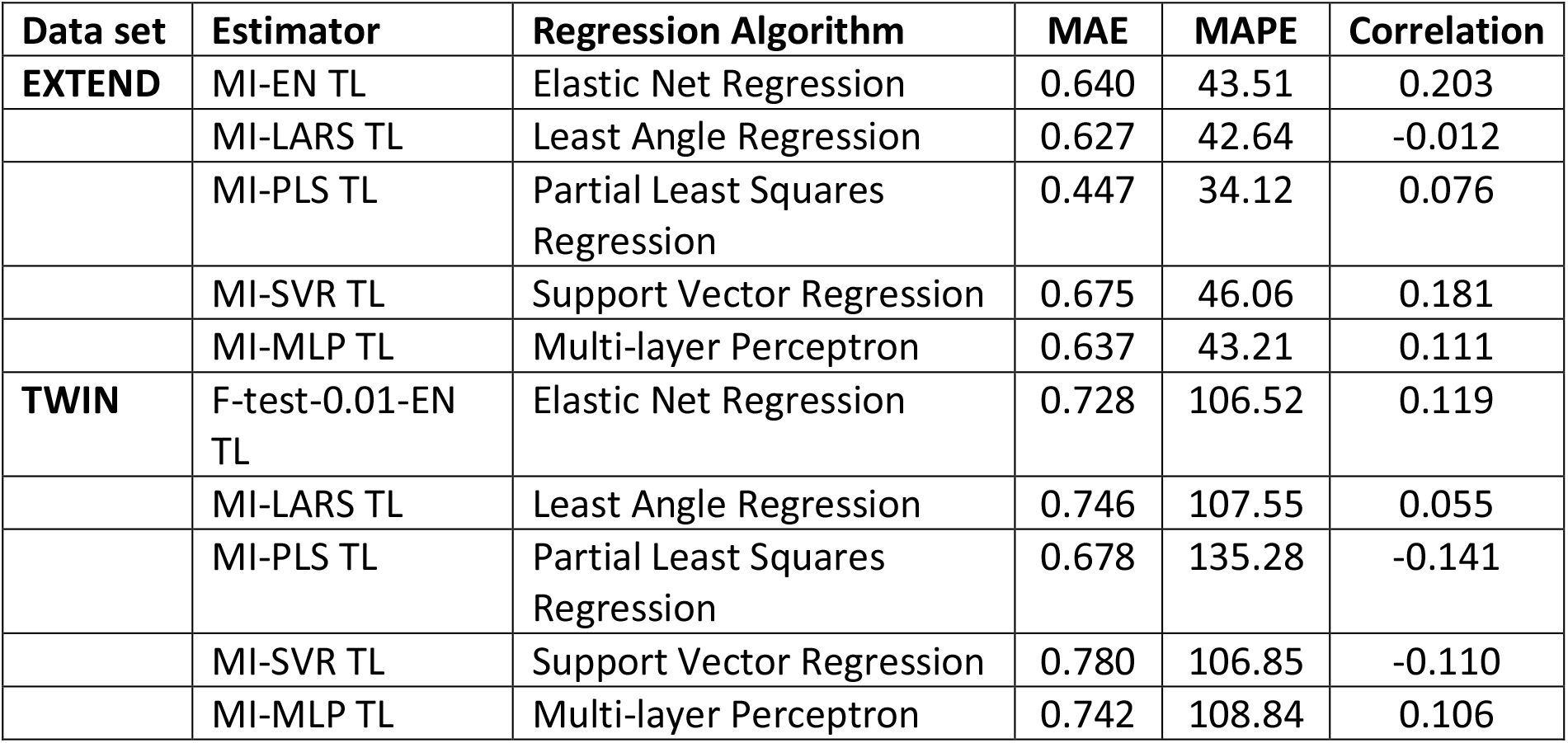
Performance scores for models constructed on Dunedin data and tested on EXTEND and TWIN data. Metrics include MAE, MAPE and Pearson correlation between predicted and actual TL. These estimators utilise mutual information and F-test (0.01 FDR) as the initial feature-selection stage for the EXTEND and TWIN data respectively - followed by an array of varying regression algorithms. For comparison the performances of MI-EN TL and F-test-0.01-EN TL is included for the EXTEND and TWIN cases respectively.

| Data set | Estimator | Regression Algorithm | MAE | MAPE | Correlation |
| --- | --- | --- | --- | --- | --- |
| EXTEND | MI-EN TL | Elastic Net Regression | 0.640 | 43.51 | 0.203 |
|  | MI-LARS TL | Least Angle Regression | 0.627 | 42.64 | -0.012 |
|  | MI-PLS TL | Partial Least Squares Regression | 0.447 | 34.12 | 0.076 |
|  | MI-SVR TL | Support Vector Regression | 0.675 | 46.06 | 0.181 |
|  | MI-MLP TL | Multi-layer Perceptron | 0.637 | 43.21 | 0.111 |
| TWIN | F-test-0.01-EN TL | Elastic Net Regression | 0.728 | 106.52 | 0.119 |
|  | MI-LARS TL | Least Angle Regression | 0.746 | 107.55 | 0.055 |
|  | MI-PLS TL | Partial Least Squares Regression | 0.678 | 135.28 | -0.141 |
|  | MI-SVR TL | Support Vector Regression | 0.780 | 106.85 | -0.110 |
|  | MI-MLP TL | Multi-layer Perceptron | 0.742 | 108.84 | 0.106 |

### 3.4 Comparison with previously developed DNAmTL estimator

The novel 140-CpG DNA methylation-based TL estimator described previously by Lu et al. [28], (DNAmTL), is compared with the range of estimators developed in our study. We utilised both validation sets (EXTEND and TWIN) for this purpose. Predictions (estimates) from each estimator were assessed via their correlation with the actual values of TL in both data sets (Table 6) - an analogous correlation analysis was conducted in [28, 67]. On the EXTEND data, the best performing estimator is that which uses PCA in advance of elastic net regression (PCA-EN TL) achieving a moderate correlation coefficient of approximately 0.3 between predicted and actual TL. The next best performing model was our mutual information with elastic net (MI-EN TL) estimator which achieved a correlation of 0.203. Comparing with the DNAmTL estimator of Lu et al. [28], this achieved a correlation of 0.216 (114 of the 140 DNAmTL CpG measures were available in EXTEND). There is substantial variation observed in the correlations reported across the estimators (Table 6), suggesting that the choice of feature-selection method is critical, given the specific nature of individual data sets.

Previously, the DNAmTL estimator developed by Lu et al. [28] was tested on a range of independent data sets. Four of these data sets contained TL measured using the gold standard Southern blot telomere restriction fragment method (TRF), with the DNAmTL estimator achieving correlations ranging r=0.41-0.5. However, on 3 data sets with quantitative polymerase chain reaction (qPCR) TL measures (i.e., similar to the TL measurement method used in our study), the correlations were observed to be variable (r=-0.01, 0.08 and 0.38). On applying the DNAmTL estimator to the Dunedin data set, a correlation of 0.242 between predicted and actual TL values was found (with 115 of 140 DNAmTL CpGs available in Dunedin data). Additionally, the DNAmTL estimator achieves correlations of 0.216 and 0.092 on the EXTEND and TWIN data sets respectively.

The observed differences in the correlations between predicted and actual TL for TRF and qPCR data, and the substantial variation in the case of qPCR may be due to known lab-to-lab variation of qPCR TL assays and/or be due to assay reproducibility [67]. The qPCR method is frequently used to measure TL due to its high-throughput and small DNA requirements; however, due to its sensitivity to pre-analytic factors such as DNA extraction or storage, its reliability is limited [72]. This suggests that potentially DNA methylation-based estimators are not as accurate for qPCR measured TL prediction. On the whole, performance of the estimators was substantially lower when tested on the TWIN data set with the estimator that applied the F-test (0.01 FDR) in conjunction with elastic net achieving the highest correlation (r=0.119) between predicted and actual TL. Comparatively, the DNAmTL estimator yielded a correlation of 0.092 on the TWIN data set, with all 140 CpG sites being available for use in the estimator.

We compared the features selected by Lu et al.’s DNAmTL estimator [28] and our MI-EN TL estimator, both of which yielded relatively small feature sets (140 and 407 features respectively). Only two features (CpGs) were found to be common to both estimators. This highlights the challenge of finding a DNA methylation-based estimator that would generalise well on all data sets. Theoretically, to acquire such a level of generalisation, one would require very large data sets, which at the present time is difficult to obtain. However, in the future, aggregation of data sources may be possible which could yield a highly generalisable estimator, wherein applying a robust methodology such as that demonstrated in this paper, should further enhance the ultimate estimator.

### 3.5 Correlation of actual TL and estimators MI-EN TL and PCA-EN TL

Plots of actual TL versus two of our best performing DNA methylation-based estimators (MI-EN TL and PCA-EN-TL) are shown in Figure 6 for the Dunedin, EXTEND and TWIN data sets. As expected, higher correlations are observed when the estimators are applied to the Dunedin training data (r=0.487 and r=0.498) with the highest correlation in the validation cohorts given by the PCA-EN TL estimator applied to the EXTEND data (r=0.295).

**Figure 6:**
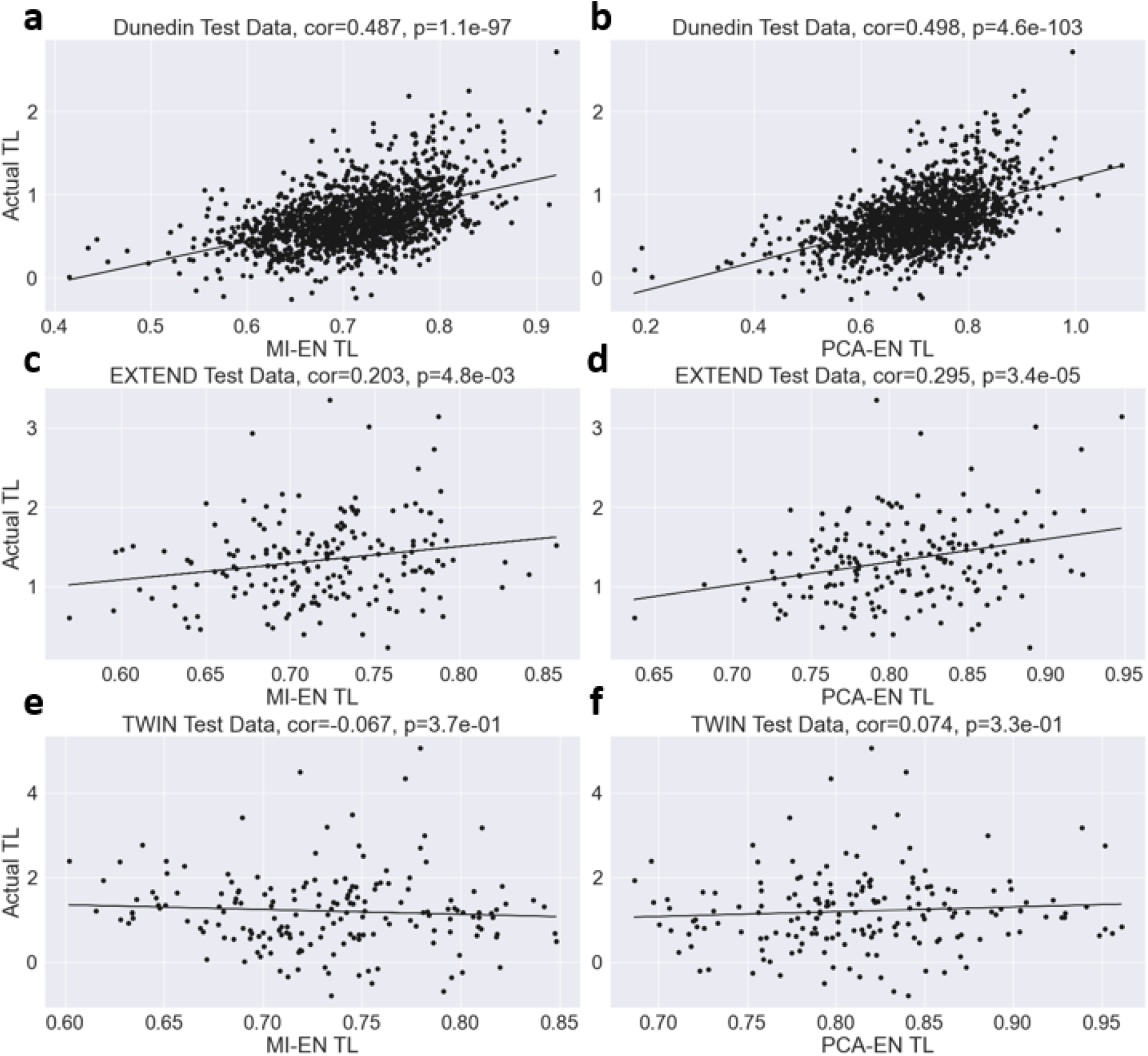
Measured relative TL versus MI-EN TL/PCA-EN TL in training and test datasets. Scatter plots of DNA methylation-based telomere length (MI-EN TL/PCA-EN TL, x-axis) versus TL measured by qPCR (y-axis). **A**. Dunedin training data (MI-EN-TL). **B**. Dunedin training data (PCA-EN-TL). **C**. Test data (EXTEND) with MI-EN TL. **D**. Test data (EXTEND) with PCA-EN TL. **E**. Test data (TWIN) with MI-EN TL. **F**. Test data (TWIN) with PCA-EN TL. As evidenced in other TL estimation studies [28, 67], it is notable that our estimators yield a more restrictive range of TL values relative to actual TL. Each panel includes a Pearson correlation coefficient and correlation test p-value.

Given the poor performance of the developed estimators on the TWIN data, no further follow-up analysis using this data set was performed.

### 3.6 Predicted TL from MI-EN TL and PCA-EN TL estimators more strongly correlated with age than TL

In the EXTEND validation data set MI-EN TL and PCA-EN TL showed stronger negative correlations with chronological age (r=-0.506 and r=-0.565 respectively) than did actual TL (r=-0.218), as shown in Figure S1. Next, a multiple regression model was constructed using the EXTEND data to explore the relationship between age and TL. Our analysis showed that the MI-EN TL and PCA-EN TL estimators, were associated with much more significant p-values (p=8.49e-14 and p=6.91e-17 respectively) than actual relative TL (p=2.83e-03), after adjusting for sex, current smoking status and other confounders (Table S1, S.I). Our findings are consistent with the Lu et al. [28] DNAmTL estimator, which achieved more significant associations with age than actual TL did with age.

Previously, an epigenetic clock, DNAmAge, was developed [5] in the form of a multi-tissue predictor of age that utilises DNA methylation data. We examined age acceleration, generated by regressing DNAmAge on age, and plotted it against actual and estimated TL for comparison (Figure S2, S.I.). We observed no correlation (r=-0.029, p=0.69) between actual TL and the age acceleration measure in EXTEND data, as shown previously ([42, 73-75]). Stronger, but small, negative correlations were observed between age acceleration and both our TL estimators MI-EN TL (r=-0.143, p=0.048) and PCA-EN TL (r=-0.161, p=0.026). Similarly, Lu et al. [28] reported a Pearson correlation of r=-0.2 between the DNAmAge age acceleration measure and their age-adjusted TL estimator, DNAmLTLadjAge.

### 3.7 Effect of sex

Previously studies have shown that women have longer TL than men, when considering groups of the same age [76]. All three of the multiple regression models outlined in Table S1 in the supplementary information indicate that age-adjusted relative TL, age-adjusted MI-EN relative TL and age-adjusted PCA-EN relative TL were longer for women than men. The p-value for age-adjusted relative TL (p=4.58e-11, n=192) was much less significant than those for age-adjusted MI-EN relative TL and age-adjusted PCA-EN relative TL (p=1.29e-94 and p=5.46e-101 respectively). Comparatively, Lu et al. [28] reported that age-adjusted TL and age-adjusted DNAmTL were longer in females than males with the p-value for age-adjusted LTL (p=2.15E-04) being much less significant than that for age-adjusted DNAmTL (p=1.14E-15).

### 3.8 MI-EN TL and PCA-EN TL association with imputed blood cell composition

The following imputed blood cell counts were analysed for the validation cohort EXTEND (B cells, naïve CD4+ T, naïve CD8+ T, exhausted cytotoxic CD8+ T cells (CD8 positive, CD28 negative, CD45R negative), plasma blasts, natural killer (NK) cells, monocytes, and granulocytes). The blood cell composition imputation of B cell, NK cells, monocytes and granulocytes were imputed using the Houseman method [77], while remaining cells were imputed based on the Horvath method [78] as described previously [42].

The abundance of naïve CD8+ T cells and memory CD8+ T cells has previously been shown to correlate with actual TL [79], and a previously developed estimator (DNAmTL) was found to be significantly correlated with several imputed measures of leucocytes e.g. naïve CD8+ T cells (r=0.42, p=2.2E-151) [28]. We found similar correlations for naïve CD8+ T cells in the EXTEND cohort (r=0.39, p=1.5E-08 and r=0.27, p=1.8E-04) for PCA-EN TL and MI-EN TL respectively. Additionally, MI-EN TL was significantly correlated with several other imputed blood cell measures i.e., CD8pCD28nCD45RAn cells (r=-0.17, p=1.8E-02), CD4T cells (r=0.19, p=8.1E-03) and natural killer cells (r=-0.16, p=2.7E-02). Similarly, PCA-EN TL was significantly correlated with CD8pCD28nCD45RAn cells (r=-0.16, p=2.6E-02), plasma blasts (r=0.196, p=6.6E-03), natural killer cells (r=-0.16, p=2.7E-02), monocytes (r=-0.21, p=4.0E-03) and granulocytes (r=0.26, p=2.7E-04). Both MI-EN TL and PCA-EN TL generally exhibited considerably stronger correlations with imputed blood cell composition than actual TL (Tables S2, S3, S4 in S.I.).

### 3.9 MI-EN TL and PCA-EN TL association with actual blood cell composition in Dunedin cohort

Actual blood cell counts (as opposed to imputed) were available for the Dunedin study. Moderate negative correlations were observed for basophil count, with the estimated telomere lengths from MI-EN TL and PCA-EN TL achieving the highest correlations (r=-0.41, p=1.8E-05 and r=-0.38, p=6.7E-05 respectively), compared to r=-0.24 (p=0.014) for the actual TL. In the case of eosinophils, actual TL showed similar negative correlation (r=-0.15, p=9.3E-5) to both PCA-EN TL and MI-EN TL (r=-0.19, p=3.6E-07 and r=-0.10, p=1.2E-02 respectively).

There was a significant small correlation of r=-0.09 (p=0.013) between actual monocyte count and measured TL for the Dunedin data, while stronger and more significant negative correlations were observed with both MI-EN TL and PCA-EN TL (r=-0.15, p=5.4E-05 and r=-0.16, p=3.1E-05). Imputed monocyte blood cell counts are available for the EXTEND cohort for comparison. Of the correlations between imputed monocyte count and actual TL, MI-EN TL and PCA-EN TL, PCA-EN TL showed the strongest correlation (r=-0.21, p=4.0E-03) with imputed monocyte count. Figure S3 (S.I.) shows plots of monocyte counts versus actual TL, MI-EN TL and PCA-EN TL for both the Dunedin and EXTEND data sets. Blood cell composition correlation details for EXTEND and Dunedin data can be viewed in Tables S2, S3, S4 and S5 in the supplementary information.

It is interesting to note that despite the Lu at al. DNAmTL estimator [28] and our MI-EN TL estimator having only two features in common, they are both shown to correlate with many of the same biological entities i.e., blood cell composition, age and sex. This suggests that our developed estimators may have captured these biological properties by finding features (CpGs) that identify them best in our data, whereas a predominantly different signature may achieve this in another data set.

## Conclusions

In summary, we have outlined a robust methodology that utilises feature-selection approaches and ML algorithms in the development of a DNA methylation-based TL estimator. We have shown through results on independent data that differences in the efficacy of developed estimators exist, primarily due to the inherently varying combination of methods and algorithms with relatively small heterogenous data sets. Consistent with research in ML and deep learning, until truly representative big data is available, it will be challenging to develop estimators of biological traits, such as TL, that generalise well to new unseen data. Interestingly, different estimators with extensively different feature sets, correlate with many of the same biological properties, which suggests that an estimator has a choice of CpGs with which to represent traits.

An interesting outcome of our work is that PCA, a technique not traditionally used for feature reduction of high dimensional data in DNA methylation studies, performs well in comparison to a range of more typically utilised approaches. This suggests that it may have utility in the development of DNA methylation-based estimators or classifiers where prediction is paramount, with identification of underlying features (CpG sites) not of primary interest. Such estimators, however, may have less clinical utility for the development of DNA methylation-based biomarkers of disease. Furthermore, the methodology adopted herein, that compares and assesses candidate estimators of TL, could easily be applied when developing estimators of other biological markers and disease phenotypes, to examine their relationship with DNA methylation and potentially improve their predictive value.

## Acknowledgments

We thank the Dunedin Study members, unit research staff, and study founder Phil Silva. We are grateful to all participants of EXETER 10000. We would like to thank the Irish Centre for High End Computing (https://www.ichec.ie/) for the use of their HPC infrastructure and support, and the Applied Intelligence Research Centre (Technological University Dublin) for provision of computing cluster resources.

## Abbreviations

CV: Cross-validation
CpG: Cytosine-Phosphate-Guanine
DNA: Deoxyribonucleic Acid
FDR: False Discovery Rate
MAE: Mean Absolute Error
MAPE: Mean Absolute Percentage Error
ML: Machine Learning
PCA: Principal Component Analysis
qPCR: Quantitative Polymerase Chain Reaction
SVR: Support Vector Regression
TL: Telomere Length
TRF: Telomere Restriction Fragment

## Availability of data and materials

Source code and scripts are available upon request. The Dunedin Study datasets reported in the current article are not publicly available due to a lack of informed consent and ethical approval for public data sharing. The Dunedin study datasets are available on request by qualified scientists. Requests require a concept paper describing the purpose of data access, ethical approval at the applicant’s university and provision for secure data access (https://moffittcaspi.trinity.duke.edu/research-topics/dunedin). Secure access offered on the Duke, Otago and King’s College campuses. For the TWIN study, data is freely available in the supplemental files of the previously published article [43]. The EXTEND study data is deposited in the Gene Expression Omnibus (GEO) database (accession number: GSE113725).

## Ethics approval and consent to participate

All experiments were performed in accordance with national guidelines and regulations as well as the Declaration of Helsinki. The Dunedin study participants gave written informed consent, and study protocols were approved by the NZ-HDEC (New Zealand Health and Disability Ethics Committee). For both EXTEND and TWIN data sets, informed consents were obtained from participants of these studies and ethical approval was obtained from University of Exeter Medical School Research Ethics Board.

## Consent for publication

## Authors’ Contributions

TMM conceived the study and with SJD supervised the project. TD performed all statistical and machine learning analysis. TD, SJD and TMM contributed to the computational interpretation of the results and were involved in the overall manuscript conceptualisation. TMM, JM, ED, EH, AC, TEM and BW provided telomere length and/or DNA methylation data for the study. AC, TEM and RP conceptualised and designed the Dunedin longitudinal cohort study. RP, DC, KS, BW, AC and TEM conceptualised data collection protocols and created variables for the Dunedin Cohort. TD, TMM and SJD drafted the manuscript. All authors reviewed and approved the final manuscript.

