## Supplementary Information for "A comparison of feature selection methodologies and learning algorithms in the development of a DNA methylation-based telomere length estimator"

#### Section 1:

##### Performance Metrics

A range of performance metrics which are commonly used in DNA methylation-based regression studies (such as age estimation) are reported in this study. These include the Mean Absolute Error, the Mean Absolute Percentage Error (MAPE), Root Mean Squared Error (RMSE) and Pearson's correlation coefficient (r).

The Mean Absolute Error (MAE), Mean Absolute Percentage Error (MAPE), and Pearson's correlation coefficient ($r_{xy}$) can be represented as:

$$MAE = n^{-1} \sum_{i=1}^{n} |x_i - x_i'| \quad (1)$$

$$MAPE = \frac{100}{n} \sum_{i=1}^{n} \left| \frac{x_i - x_i'}{x_i} \right| \quad (2)$$

$$r_{xy} = \frac{n \sum X_i Y_i - \sum X_i \sum Y_i}{\sqrt{n \sum X_i^2 - (\sum X_i)^2} \sqrt{n \sum Y_i^2 - (\sum Y_i)^2}} \quad (3)$$

where $x_i$ and $x_i'$ are the actual and model predicted values for sample $i$ respectively, $n$ is the number of samples, $X_i$ and $Y_i$ are a pair of random variables [1].

### Supplementary Figures.

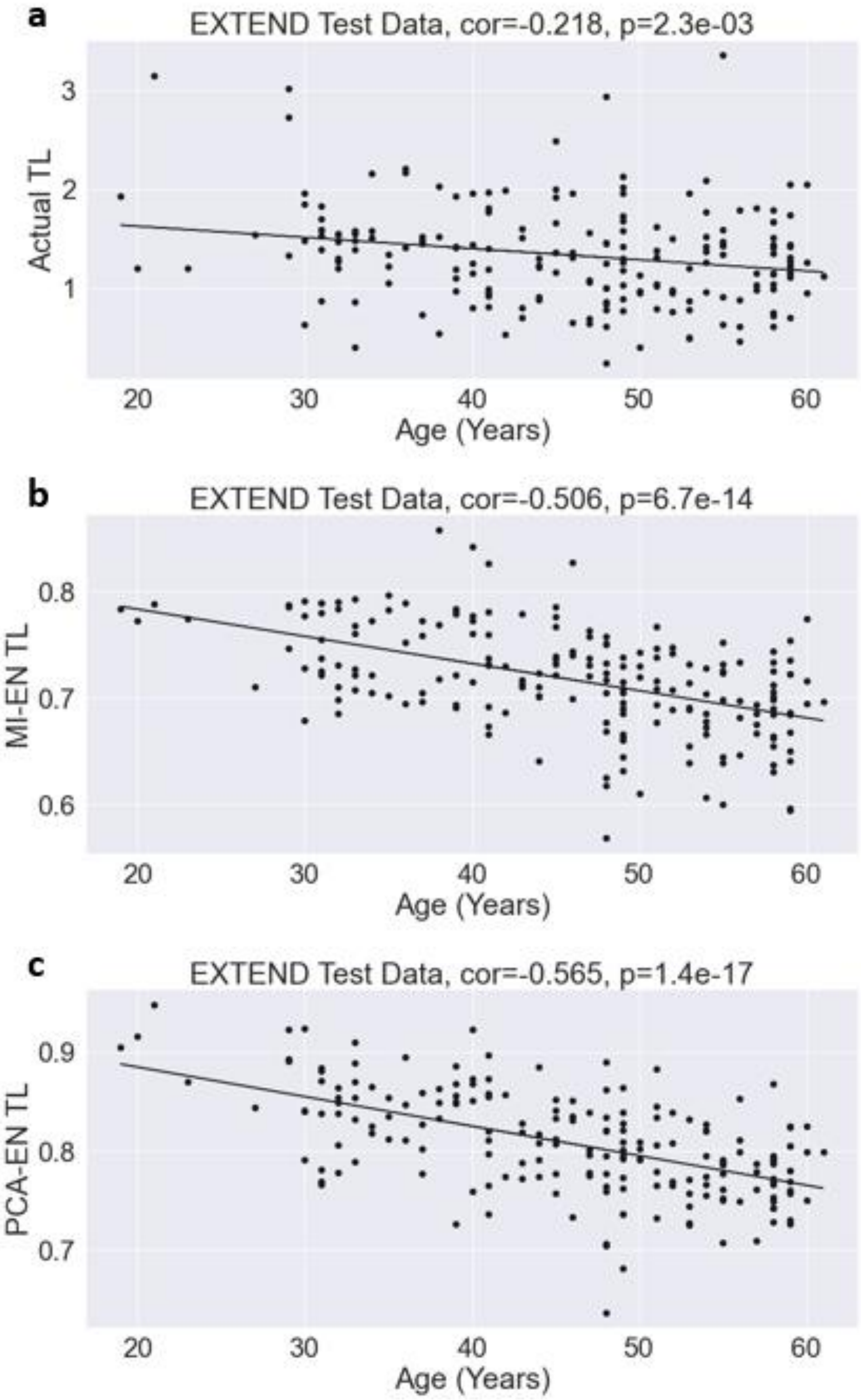


Figure S1: **A**. Chronological age vs. measured TL, **B**. Chronological age vs. MI-EN TL and **C**. Chronological age vs. PCA-EN TL. Plots relate to the EXTEND test data set and Pearson's correlation coefficient and correlation test p-value is reported for each case.

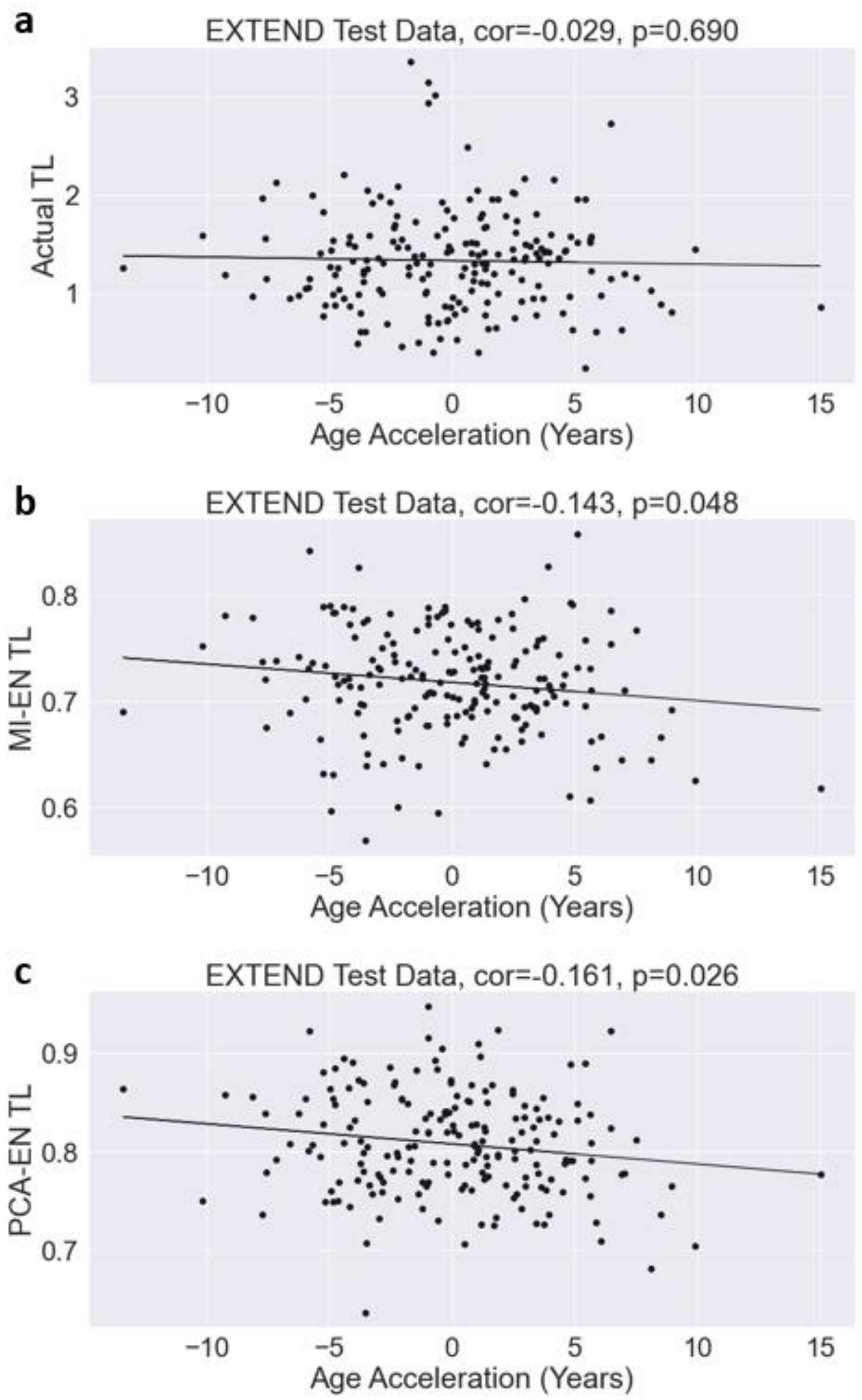


Figure S2: The x-axis variable is Age Acceleration, which represents the residual of the Lu et al. estimator DNAmAge [2] regressed on age. **A.** Age Acceleration vs. measured TL, **B.** Age Acceleration vs. MI-EN TL and **C**. Age Acceleration vs. PCA-EN TL. Plots relate to the EXTEND test data set and Pearson's correlation coefficient and correlation test p-value is reported for each case.

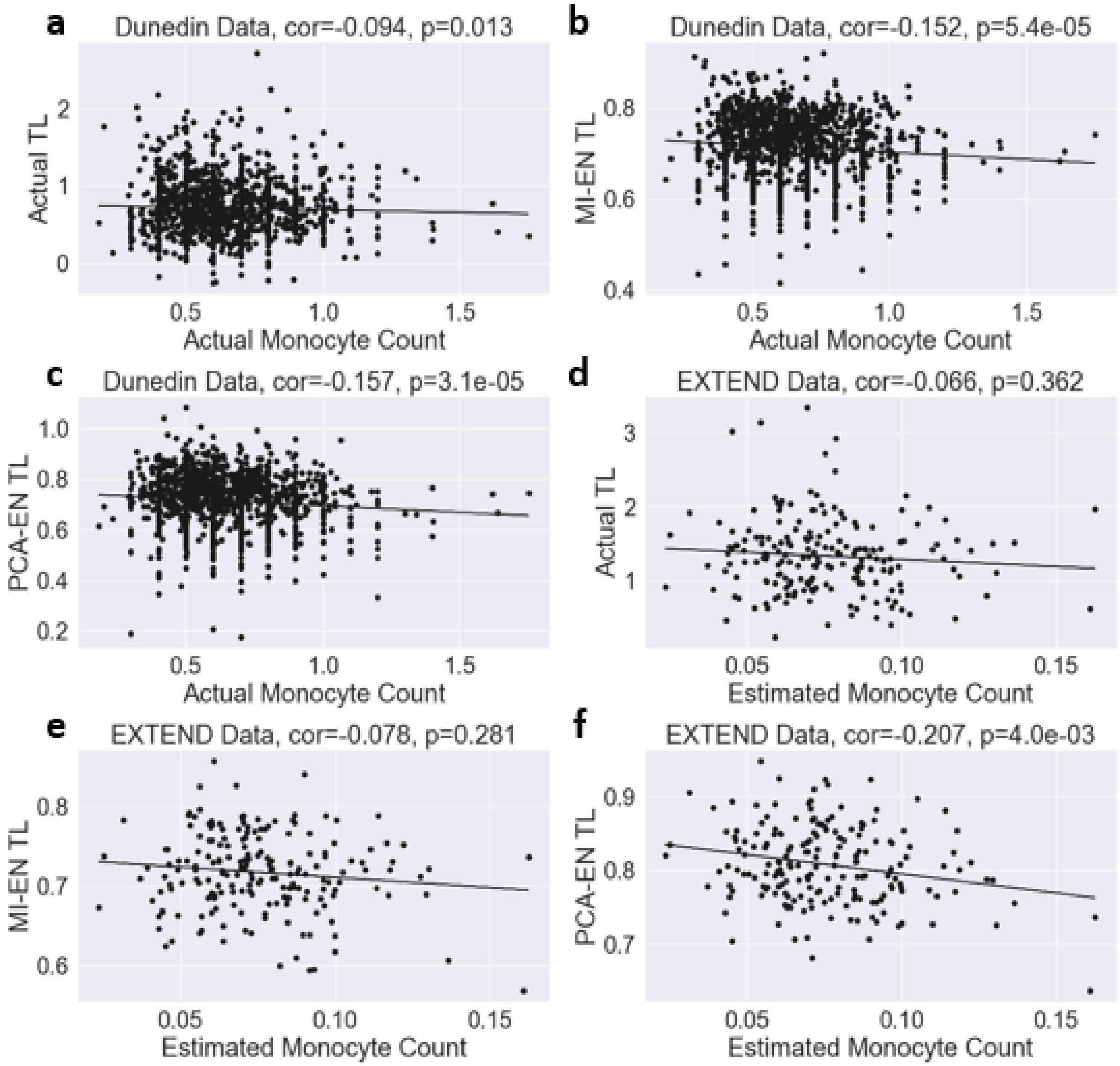


Figure S3: **A.** Actual Monocyte Count vs. measured TL, **B.** Actual Monocyte Count vs. MI-EN TL, **C**. Actual Monocyte Count vs. PCA-EN TL, **D.** Estimated Monocyte Count vs. measured TL, **E.** Estimated Monocyte Count vs. MI-EN TL, **F**. Estimated Monocyte Count vs. PCA-EN TL Dunedin data contained actual monocyte counts, EXTEND data set contained estimated monocyte counts. Pearson's correlation coefficient and correlation test p-value is reported for EXTEND data while repeated measures correlation was used for Dunedin data due to many donors having multiple samples.

### Supplementary Tables.

Table S1: Multiple linear regression for analysis of biological correlates in the EXTEND data set. The top panel contains results from a multiple linear regression model analysis of actual TL (dependent variable) on a range of covariates for the EXTEND data set (n=192). The model was regressed on age, sex, current smoking status and body mass index (BMI). The middle and bottom panels contain results of analogous multiple regression models but with dependent variables of MI-EN TL and PCA-EN TL respectively. SE denotes the standard error.

| **Actual TL values** | | | |
|---|---|---|---|
| **Variable** | **Coefficient (SE)** | **t-statistic** | **P-value** |
| Intercept | 1.042 (0.170) | 6.13 | 5.13e-09 |
| Age | -0.011 (0.004) | -3.03 | 2.83e-03 |
| Female | 0.605 (0.086) | 7.00 | 4.58e-11 |
| Male | 0.437 (0.102) | 4.28 | 2.95e-05 |
| Current Smoker | 0.003 (0.183) | 0.02 | 0.988 |
| BMI | 0.009 (0.008) | 1.20 | 0.232 |
| **MI-EN Predicted TL** | | | |
| Intercept | 0.530 (0.014) | 38.68 | 3.73e-91 |
| Age | -0.002 (0.000) | -8.07 | 8.49e-14 |
| Female | 0.283 (0.007) | 40.57 | 1.29e-94 |
| Male | 0.248(0.008) | 30.05 | 1.64e-73 |
| Current Smoker | 0.015 (0.015) | 0.99 | 0.325 |
| BMI | 0.001 (0.001) | 1.46 | 0.14 |
| **PCA-EN Predicted TL** | | | |
| Intercept | 0.625 (0.014) | 43.38 | 1.54e-99 |
| Age | -0.003 (0.000) | -9.20 | 6.91e-17 |
| Female | 0.324 (0.007) | 44.24 | 5.46e-101 |
| Male | 0.301 (0.009) | 34.77 | 1.45e-83 |
| Current Smoker | -0.008 (0.015) | -0.50 | 0.619 |
| BMI | -5.87e-05 (0.001) | -0.09 | 0.929 |

Table S2: Pearson correlation coefficient and p-values for age-adjusted actual TL and imputed blood cell counts.

| data | var | cell | Correlation | P |
|---|---|---|---|---|
| EXTEND | TLadjAge | CD8.naive | 0.120 | 9.77E-02 |
| EXTEND | TLadjAge | CD8pCD28nCD45RAn | -0.046 | 5.24E-01 |
| EXTEND | TLadjAge | Plasma blast | 0.033 | 6.53E-01 |
| EXTEND | TLadjAge | CD4T | -0.153 | 3.42E-02 |
| EXTEND | TLadjAge | NK | -0.186 | 9.73E-03 |
| EXTEND | TLadjAge | Mono | -0.066 | 3.62E-01 |
| EXTEND | TLadjAge | Gran | 0.205 | 4.28E-03 |

Table S3: Pearson correlation coefficient and p-values for age-adjusted MI-EN TL and imputed blood cell counts.

| data | var | cell | Correlation | P |
|---|---|---|---|---|
| EXTEND | MI-ENadjAge | CD8.naive | 0.268 | 1.75E-04 |
| EXTEND | MI-ENadjAge | CD8pCD28nCD45RAn | -0.171 | 1.79E-02 |
| EXTEND | MI-ENadjAge | Plasma blast | -0.099 | 1.74E-01 |
| EXTEND | MI-ENadjAge | CD4T | 0.191 | 8.11E-03 |
| EXTEND | MI-ENadjAge | NK | -0.160 | 2.69E-02 |
| EXTEND | MI-ENadjAge | Mono | -0.078 | 2.81E-01 |
| EXTEND | MI-ENadjAge | Gran | -0.039 | 5.92E-01 |

Table S4: Pearson correlation coefficient and p-values for age-adjusted PCA-EN TL and imputed blood cell counts.

| data | var | cell | Correlation | P |
|---|---|---|---|---|
| EXTEND | PCA-ENadjAge | CD8.naive | 0.395 | 1.49E-08 |
| EXTEND | PCA-ENadjAge | CD8pCD28nCD45RAn | -0.160 | 2.63E-02 |
| EXTEND | PCA-ENadjAge | Plasma blast | 0.196 | 6.55E-03 |
| EXTEND | PCA-ENadjAge | CD4T | -0.011 | 8.85E-01 |
| EXTEND | PCA-ENadjAge | NK | -0.159 | 2.73E-02 |
| EXTEND | PCA-ENadjAge | Mono | -0.207 | 4.02E-03 |
| EXTEND | PCA-ENadjAge | Gran | 0.260 | 2.72E-04 |

Table S5: Repeated measures correlation between actual TL/MI-EN TL/PCA-EN TL and actual blood cell counts.

| data | var | cell | Correlation | P |
|---|---|---|---|---|
| Dunedin | Actual TL | Neutrophils | -0.035 | 0.34082800 |
| Dunedin | Actual TL | Lymphocytes | -0.036 | 0.33419100 |
| Dunedin | Actual TL | Monocytes | -0.094 | 0.01263800 |
| Dunedin | Actual TL | Eosinophils | -0.150 | 0.00009300 |
| Dunedin | Actual TL | Basophils | -0.242 | 0.01359000 |
| Dunedin | MI-EN TL | Neutrophils | -0.054 | 0.14370800 |
| Dunedin | MI-EN TL | Lymphocytes | -0.135 | 0.00026500 |
| Dunedin | MI-EN TL | Monocytes | -0.152 | 0.00005400 |
| Dunedin | MI-EN TL | Eosinophils | -0.096 | 0.01210500 |
| Dunedin | MI-EN TL | Basophils | -0.408 | 0.00001800 |
| Dunedin | PCA-EN TL | Neutrophils | -0.021 | 0.57911400 |
| Dunedin | PCA-EN TL | Lymphocytes | -0.134 | 0.00029700 |
| Dunedin | PCA-EN TL | Monocytes | -0.157 | 0.00003100 |
| Dunedin | PCA-EN TL | Eosinophils | -0.194 | 0.00000036 |
| Dunedin | PCA-EN TL | Basophils | -0.382 | 0.00006700 |
